# Identification of a neural basis for energy expenditure in the arcuate hypothalamus

**DOI:** 10.1101/2025.03.11.642601

**Authors:** Ting Wang, Shuping Han, Yaxin Wang, Yuxiao Li, Shuangfeng Zhang, Feipeng Zhu, Zhen-Hua Chen, Zhifang Xing, Yu Zheng Li, Jingjing Wang, Mingrui Xu, Qinghua Liu, Man Jiang, Xiaohong Xu, Xiangning Li, Peng Cao, Hui Gong, Qing-Feng Wu

**Affiliations:** State Key Laboratory of Molecular Developmental Biology, Institute of Genetics and Developmental Biology, Chinese Academy of Sciences, Beijing 100101, China; University of Chinese Academy of Sciences, Beijing 100101, China; State Key Laboratory of Digital Medical Engineering, School of Biomedical Engineering, Hainan University, Haikou 570228, China; National Institute of Biological Sciences, Beijing 102206, China; Department of Physiology, School of Basic Medicine and Tongji Medical College, Huazhong University of Science and Technology, Wuhan 430030, China; Institute of Neuroscience, State Key Laboratory of Neuroscience, CAS Center for Excellence in Brain Science and Intelligence Technology, Chinese Academy of Sciences, Shanghai 200031, China; HUST-Suzhou Institute for Brainsmatics, JITRI, Suzhou 215123, China; Beijing Key Laboratory for Genetics of Birth Defects, Beijing 100045, China

## Abstract

Given the evolutionary instinct for caloric intake and the tendency for weight rebound after discontinuing dietary interventions or medications, increasing energy expenditure emerges as an alternative obesity treatment. However, neural regulation of energy expenditure remains poorly understood. Here, we report that a hypothalamic neuronal subtype, characterized by *Crabp1* expression, establishes connections with multiple hypothalamic nuclei to regulate energy expenditure in mice. Inactivation of Crabp1 neurons reduces physical activity, body temperature, and adaptive thermogenesis, leading to an obese phenotype. Conversely, activation of these neurons increases energy expenditure and mitigates diet-induced obesity. Structural and functional analyses reveal that Crabp1 neurons promote energy metabolism through a “one-to-many” projection pattern. While Crabp1 neurons are rapidly activated by cold exposure and physical activity, prolonged light exposure abrogates their firing and may mediate light-induced metabolic disorder. Together, we reveal a neural basis that integrate various physiological and environmental stimuli to control energy expenditure and body weight.

## Introduction

With the escalating prevalence of obesity, there is a pressing need to elucidate the regulatory mechanisms underlying body weight control and develop novel strategies for therapeutic intervention.^1,2^ Research over recent decades has established the hypothalamus, particularly the arcuate nucleus (ARC), as a central hub for homeostatic control of energy balance.^3,4^ An excess of energy intake over expenditure results in positive energy balance, leading to weight gain and body fat storage. Established model of energy homeostasis has spotlighted a yin-yang interplay between two well-defined neuronal subtypes in the hypothalamic arcuate nucleus: proopiomelanocortin (POMC) neurons and agouti-related neuropeptide (AgRP) neurons. Activation of POMC neurons reduces food intake and increases energy expenditure, whereas AgRP neurons elicit increased food consumption and reduced energy expenditure when activated.^5–7^ Notably, the ARC hosts many other neuronal subtypes that are distinct from POMC and AgRP neurons, such as “RIP-Cre” and “Pdx1-Cre” neurons.^8,9^ These neurons serve important functions with regards to hormone release, appetite control, puberty onset, and energy homeostasis. For example, selective activation of RIP-Cre neurons enhances energy expenditure without altering food intake, whereas Pdx1-Cre neurons promote feeding behavior via GABA release during the postweaning period.^8,9^ Unfortunately, the molecular identity of “non-AgRP, non-POMC” neurons like these remains unclear. Given the intricate organization of the arcuate neurons, it is crucial to dissect the molecular, structural, and functional complexity of these transcriptionally undefined neurons in the hypothalamus.

To unravel the intricacies of ARC, a hierarchical approach of navigating the molecularly defined neuronal diversity has been proposed. This involves inspecting the major functional response types (e.g., glutamatergic and GABAergic), followed by iterative refinement among main neuronal subtypes to achieve the finest resolution along the hierarchy of transcriptional similarity.^10–12^ In the ARC, it is well-known that glutamatergic neurons activation rapidly regulates satiety, while chronic activation of GABAergic neurons induces food intake and causes obesity.^13,14^ Yet, molecular classification of the main “non-AgRP, non-POMC” neuronal subtypes and their functional diversity are not fully understood.

Cell type-specific neuronal connectivity, operating within and extending beyond the hypothalamus, forms the structural foundation for understanding neural control of energy balance. Various neuronal subtypes are known to exert distinct functions based on their unique connectivity and molecular properties. The emerging paradigm suggests that a single molecularly-defined neuronal subtype may exhibit different functions by projecting to distinct sets of downstream targets. For instance, AgRP neurons project to dorsal raphe nucleus to control energy expenditure but not food intake, while their innervation of the paraventricular hypothalamus stimulates feeding behavior.^15–17^ Thus, mapping the neural circuit structure of each neuronal subtype is instructive for comprehending its function under both physiological and pathological conditions.

From a translational standpoint, genetic identification of neural circuits governing energy intake and expenditure has been facilitating the development of treatments for obesity and associated metabolic disorders.^18,19^ Although calorie restriction via fasting or drug-induced appetite suppression effectively reduces body weight, weight rebound in the long term is a common occurrence due to synaptic amplification and epigenetic modifications within hypothalamic circuits during caloric deficit.^20,21^ Therefore, dissecting the neuronal substrates and circuits that promote increased energy expenditure without suppressing appetite may circumvent the dieting dilemma and offer an alternative strategy to control body weight.

In this study, we employed single-cell and single-nucleus RNA-sequencing to identify a neuronal subtype within the ARC characterized by expression of *Crabp1* gene encoding cellular retinoic acid binding protein 1. This population is distinct from POMC and AgRP neurons, as well as neuroendocrine neurons producing growth hormone-releasing hormone (GHRH), kisspeptin, or dopamine. We demonstrate that Crabp1 neurons, in response to ambient temperature change and exercise, bidirectionally regulate energy expenditure such as physical activity and thermoregulation. By structural and functional mapping of the circuitry, we reveal that a subset of Crabp1 neurons with predominant intra-hypothalamic projections control multiple components of energy expenditure via axon collateral branches. Interestingly, prolonged light exposure reduces the activity of Crabp1 neurons, thereby contributing to decreased energy expenditure and weight gain. Thus, we identify Crabp1 neurons as an important neuronal substrate in coordinating energy expenditure under various environmental and physiological contexts, rendering it a promising target for the prevention and treatment of obesity.

## Results

### Molecular profiling of neuronal subtypes in the arcuate hypothalamus

Single-cell transcriptomic profiling has recently been applied to dissect cell types and cell states in the ARC,^11,12^ providing valuable insights into its cellular diversity. However, inconsistencies arise in the classification of neuronal subtypes across single-cell datasets generated by different technologies, even within a single hypothalamic nucleus, which complicates our understanding of neuronal diversity in the brain. To reconcile the existing inconsistency, we adopted two distinct strategies to establish a coherent classification. Building upon our recent findings that Tbx3 specifies ARC and Tbx3-derived cell lineage predominantly populates this nucleus,^22^ we first sorted cells from Tbx3 lineage at juvenile stage and performed single-cell RNA-sequencing (scRNA-seq), enabling us to identify 9 distinct neuronal subtypes from a pool of 3,300 qualified neurons (Figures 1A and 1B). With this strategy, we revealed a specific neuronal subtype featured by the absence of *Agrp*, *Pomc*, *Tac2*, *Ghrh* and *Th* expression, while exhibiting robust *Crabp1* expression (Figure 1C). As a complementary strategy, we microdissected the ARC from adult mice for single-nucleus RNA-sequencing (snRNA-seq) and obtained a total of 52,022 cells across 8 replicates after quality control (Figure S1A and Table S1). Unsupervised clustering analysis of the subsetted neurons (n = 26,208) yielded 9 clusters largely concordant with the molecular categorization obtained from the scRNA-seq data, confirming the presence of Crabp1 neurons alongside AgRP and POMC subtypes (Figures 1D, 1E, S1B and S1C; Table S2). This finding coincides with previous studies that profiled arcuate Crabp1 neurons.^23,24^

**Figure 1.**
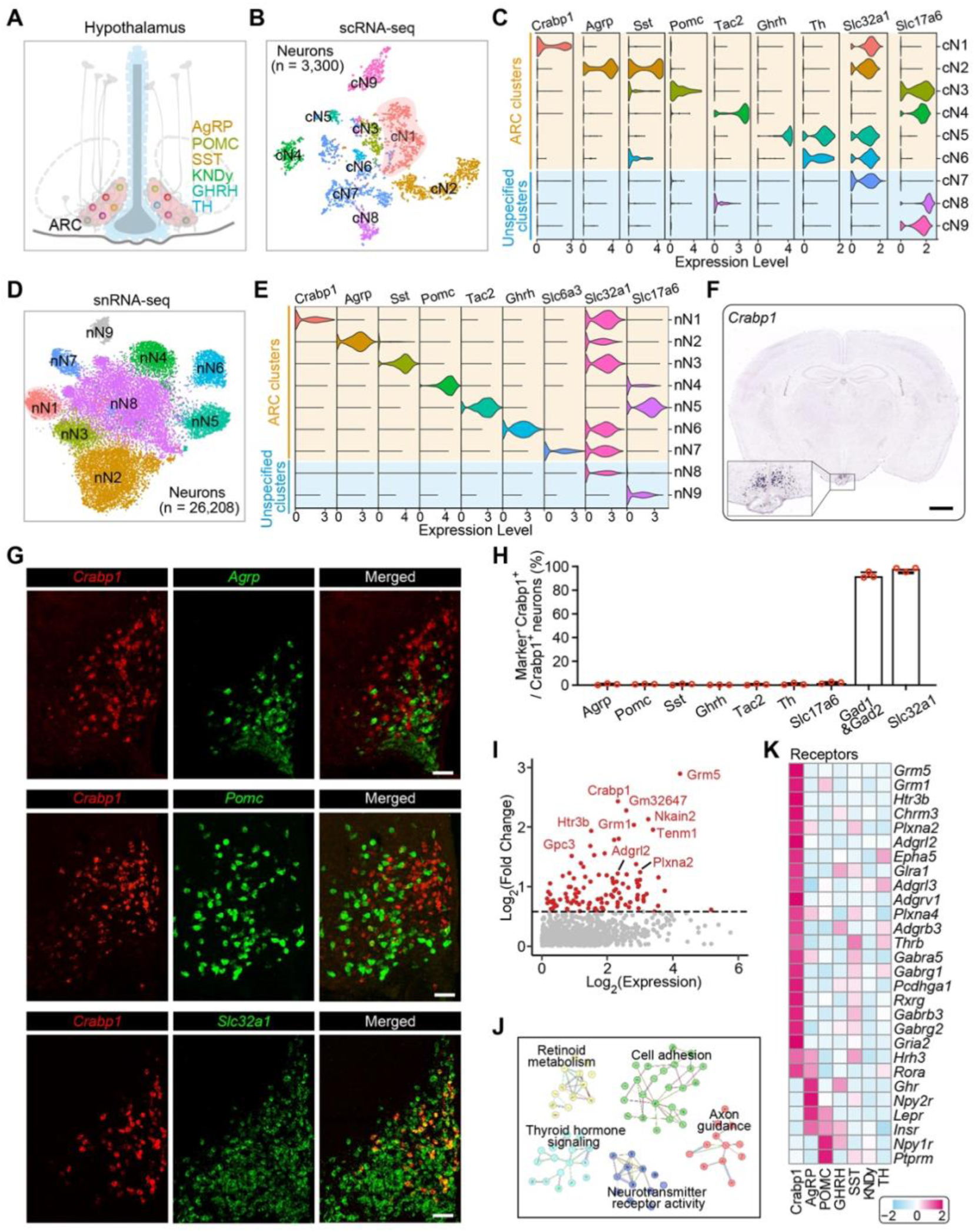
Identification of Crabp1 neurons as a population of “non-AgRP, non-POMC” neurons in the ARC. (**A**) Schematic diagram showing the heterogeneity of neuronal subtypes in the arcuate nucleus (ARC). (**B**) Unsupervised clustering analysis identifies 9 neuronal subtypes among 3,300 neurons derived from Tbx3^+^ cell lineage. We labeled Tbx3^+^ hypothalamic progenitor cells at embryonic day 9 (E9) using *Tbx3-CreER::Ai9* mice and sorted tdTomato^+^ progeny cells at postnatal day 14 (P14) for single-cell RNA sequencing (scRNA-seq). (**C**) Violin plots showing marker gene expression levels across distinct neuronal clusters, identified by scRNA-seq, within the ARC and adjacent hypothalamic nuclei. (**D**) Clustering analysis of 26,206 hypothalamic neurons from the ARC also identifies 9 neuronal subtypes. The ARC was microdissected and dissociated for single-nucleus RNA sequencing (snRNA-seq). (**E**) Violin plots indicating marker gene expression levels across diverse arcuate neuronal subtypes identified by snRNA-seq. (**F**) Image showing i*n situ* hybridization of *Crabp1* on a coronal section of adult mouse brain. The data was derived from Allen Mouse Brain Atlas. Scale bar, 1mm. (**G**) Single-molecule fluorescence *in situ* hybridization (smFISH) of *Crabp1*, *Agrp*, *Pomc*, and *Slc32a1* mRNA in the ARC indicates Crabp1 neurons as a population of “non-AgRP, non-POMC” GABAergic neurons. Scale bars, 50 μm. (**H**) Quantitative analysis of dual-color smFISH assay showing that Crabp1 neurons do not overlap with *Agrp-*, *Pomc-*, *Sst-*, *Ghrh*-, *Tac2*-, *Th*-, and *Slc17a6*-expressing cells, but co-express *Gad1*, *Gad2*, and *Slc32a1*. Values represent mean ± SEM (n = 3 brains). (**I**) Pseudobulk differential expression analysis uncovering the genes enriched in Crabp1 neurons. Genes with adjusted *P* value < 0.05 and fold change > 1.5 are highlighted in red. (**J**) Functional enrichment analysis of genes featured by Crabp1 neurons using the STRING database. (**K**) Heatmap showing the enriched expression of receptors in various neuronal subtypes, especially Crabp1 neurons.

To map the spatial distribution of Crabp1 neurons, we cross-referenced the marker with *in situ* hybridization (ISH) data from the Allen Mouse Brain Atlas (Figure 1F), and further conducted single-molecule fluorescent ISH throughout the brain. Our results corroborated that Crabp1 neurons were spatially specific to ARC, spanning its anterior, middle and posterior subregions (Figures S1D-S1F). Dual-color ISH data further demonstrated Crabp1 neurons as distinct from POMC, AgRP, SST and various neuroendocrine subtypes (Figures 1G, 1H and S1G-S1J), coinciding with our single-cell profiling data. Notably, the vast majority of Crabp1 neurons co-expressed *Slc32a1*, *Gad1* and *Gad2*, while minimal expression of *Slc17a6* was observed (Figures 1G, 1H, S1K and S1L), indicating their GABAergic identity.

To elucidate the cell type-specific transcriptional features, we performed comparative transcriptomic analysis between different neuronal subtypes using our snRNA-seq dataset, unveiling a gene set specifically enriched in Crabp1 neurons (Figures 1I, S2A and S2B; Table S3). Further network enrichment analysis identified their distinctive molecular signatures, including cell adhesion, retinoic metabolism, thyroid hormone signaling, and neurotransmitter receptor activity (Figures 1J, S2C and S2D). Unlike AgRP and POMC neuronal subtypes, Crabp1 neurons exhibited minimal expression of both insulin and leptin receptors (encoded by *Insr* and *Lepr*) but robustly expressed *Grm5*, *Htr3b*, *Glra11* and *Gabra5* encoding metabolic glutamate, serotonin, glycine and GABAergic receptors (Figures 1K, S2E and S2F). This suggests an intricate regulation of Crabp1 neuronal activity by interwoven circuits rather than circulating hormones.

### Synaptic inactivation of arcuate Crabp1 neurons reduces energy expenditure

To elucidate the functional significance and structural connectivity of Crabp1 neurons, we used CRISPR-Cas9 system to generate a knock-in mouse line expressing codon-improved Cre recombinase (iCre) driven by the endogenous *Crabp1* promoter (Figure 2A). Stereotaxic injection of adeno-associated virus (AAV) carrying EGFP in a double-inverted open (DIO) reading frame into the ARC of adult *Crabp1-iCre* mice resulted in robust genetic labeling of Crabp1 neurons, achieving ∼95% labeling specificity (Figures 2B-2D). Next, AAV-DIO-TeNT-mCherry was bilaterally injected into the ARC to disrupt synaptic transmission of Crabp1 neurons, which is widely used as a molecular tool for synaptic silencing.^25^ Four weeks post viral delivery, we observed a significant increase in body weight and fat mass of adult mice compared to control animals (Figures 2E-2G, S3A and S3B).

**Figure 2.**
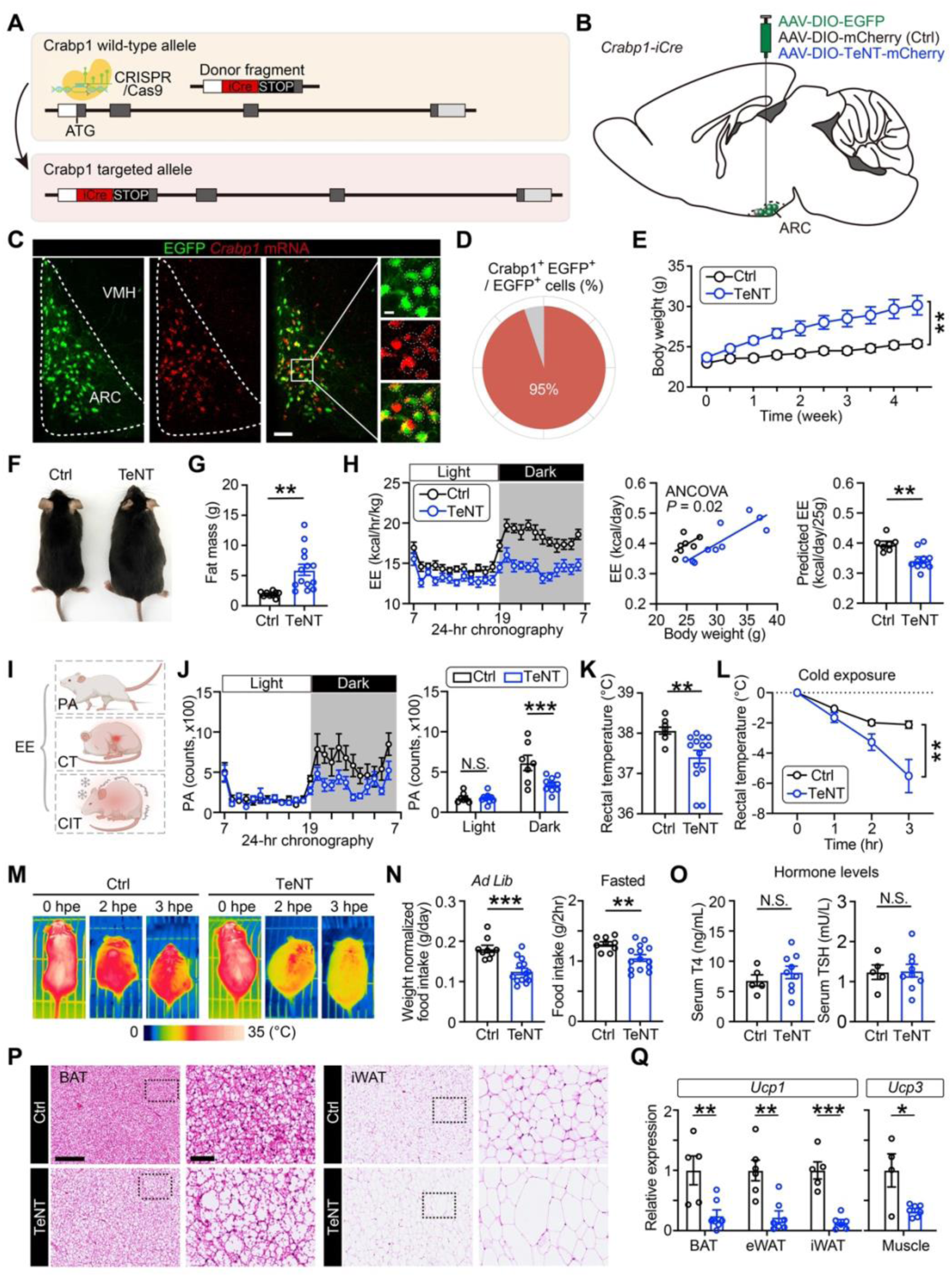
Synaptic inactivation of Crabp1 neurons suppresses energy expenditure and causes obesity. (**A**) Targeting strategy for generating *Crabp1-iCre* mouse line. (**B**) Schematic diagram showing viral injection strategy for genetic labeling and synaptic silencing of ARC Crabp1 neurons. DIO, double floxed inverted; Ctrl, control; TeNT, tetanus neurotoxin. (**C**) Representative images showing the colocalization of Cre-driven EGFP expression with the endogenous *Crabp1* mRNA puncta. Scale bars, 50 μm and 35 μm for magnified images. (**D**) Quantitative analysis of AAV-mediated genetic labeling specificity in ARC Crabp1 neurons with *Crabp1-iCre* mouse line (n = 3 brains). (**E**-**G**) Body weight dynamics (E), representative picture (F), and fat mass (G) of control and Crabp1 neuron-inactivated mice observed over the 4 weeks of monitoring (n = 9 for control group; n = 14 for TeNT group). All mice were fed with normal chow diet. (**H**) Total energy expenditure (EE; left), EE regression against body weight (middle), and predicted EE with 25g body weight (right) for control and Crabp1 neuron-silenced mice (n = 7 for control group; n = 10 for TeNT group). Mouse EE was assessed by an open flow calorimetry. Statistical analysis was performed using regression-based analysis of covariance (ANCOVA) with EE as dependent variable and body mass as covariate. (**I**) Schematic illustrating key measurable components of EE. PA, physical activity; CT, core temperature; CIT, cold-induced temperature. (**J**) Mean curves (left) and bar plots (right) showing spontaneous locomotion activity in *ad libitum* chow-fed mice during light and dark phases (n = 7 for control group; n = 10 for TeNT group). (**K**) Quantification of rectal temperature in control (n = 7) and TeNT-perturbed (n = 10) mice. (**L**) Quantitative analysis of temperature decreases in control (n = 7) and TeNT-perturbed (n = 10) during 3 hours of cold exposure at 4°C. (**M**) Representative infrared thermograph images showing the dynamics of surface temperature during 4°C cold exposure. hpe, hours post cold exposure. (**N**) Quantification of daily food intake under *ad libitum* condition (left) and 2 hours food consumption following overnight fasting (right) in control (n = 9) and TeNT-perturbed (n = 14) mice. (**O**) Quantification of serum tetraiodothyronine (T4) and thyroid-stimulating hormone (TSH) levels (n = 5 for control group; n = 9 for TeNT group). (**P**) Sample hematoxylin and eosin (H&E) staining images of brown adipose tissue (BAT) and inguinal white adipose tissue (iWAT), with magnified boxed regions highlighted. Scale bars, 250 μm and 50 μm for magnified images. (Q) Gene expression analysis of *Ucp1* levels in various adipose tissues and *Ucp3* levels in muscle (n = 5 for control group; n = 7 for TeNT group). Values are shown as mean ± SEM. Significance was analyzed by two-way analysis of variance (ANOVA) with Sidak’s multiple comparison test or unpaired two-tailed Student’s *t* test. N.S., not significant; *, *P* < 0.05; **, *P* < 0.01; ***, *P* < 0.001.

To understand how Crabp1 neurons contribute to energy balance, we further analyzed the energy expenditure and food intake in mice following neuronal silencing. Using an open flow calorimetry system, we observed that synaptic inactivation of Crabp1 neurons significantly reduced energy expenditure, oxygen consumption and carbon dioxide production in adult mice (Figures 2H and S3C-S3E). Given that energy expenditure comprises of multiple components, such as physical activity and thermoregulation,^26^ we aimed to dissect the extent to which Crabp1 neurons regulate each of these components (Figure 2I). We first analyzed the diurnal physical activity and found that genetic silencing of Crabp1 neurons reduced spontaneous locomotor activity during dark phase (Figure 2J). Second, we measured rectal temperature in light phase and found a reduction in the core temperature of mice following neuronal inactivation (Figure 2K). Last, we subjected the animals to acute cold exposure in a 4°C temperature-controlled chamber and monitored their body temperature with both infrared camera and thermometer. Our results showed that there was a substantial decrease in surface and rectal temperature of mice with synaptically-silenced Crabp1 neurons during the 4 hours cold exposure, while control mice were able to maintain their core temperature (Figures 2L and 2M). Of note, synaptic silencing of Crabp1 neurons did not increase but rather decreased food intake under both *ad libitum* and fasting conditions (Figures 2N and S3F), implicating that inactivation-induced obesity is attributed to reduced energy expenditure rather than increased energy intake.

Neural circuitry and endocrine hormone circulation are established as the primary modes of communication between peripheral tissues and central nervous system for regulating energy expenditure. Given that thyroid hormone and stress hormone are considered as important humoral regulators of energy metabolism,^27,28^ we selectively silenced Crabp1 neurons in mice and analyzed their serum hormone levels. Our data demonstrated that thyroid-stimulating hormone (TSH), tetraiodothyronine (T4), cortisol (CORT), and adrenocorticotropic hormone (ACTH) levels were not changed (Figures 2O, S3G and S3H), supporting our hypothesis that Crabp1 neurons may regulate energy expenditure via neural circuitry.

Neural control of metabolism requires the coordinated actions of peripheral tissues including brown adipose tissue (BAT), white adipose tissue (WAT), muscle, and liver. Consistent with the obesity phenotype, we observed that inactivation of Crabp1 neurons led to an accumulation of unilocular lipid droplets within interscapular BAT and larger adipocyte size in both inguinal and epididymal WAT (Figures 2P, S3I and S3J). Moreover, we assessed the expression of candidate genes associated with energy expenditure and thermogenesis in multiple peripheral tissues by reverse transcription quantitative polymerase chain reaction (RT-qPCR) analysis. Ucp1 in adipose tissue and Ucp3 in muscle function to uncouple ATP production from mitochondrial respiration, thereby dissipating energy as heat and increasing energy expenditure.^29,30^ Our results showed that silencing Crabp1 neurons reduced *Ucp1* and *Ucp3* expression in adipose tissues and muscle, respectively (Figure 2Q). Additional thermogenesis-related genes (e.g., *Dio2*, *Cidea*, *Ppara,* and *Ppargc1a*) were downregulated in adipose tissues, along with a decrease in *Fgf21* expression in the liver (Figures S3K-S3O). These results collectively imply that Crabp1 neurons may constitute a critical component within the central circuit module that remotely controls multiple aspects of energy expenditure in peripheral tissues.

### Activation of Crabp1 neurons increases energy expenditure

Next, we employed a chemogenetic tool that allows the cell type-specific expression of stimulatory designer receptor exclusively activated by designer drugs (DREADD) by bilaterally injecting AAV-DIO-hM3Dq-mCherry into the ARC of *Crabp1-iCre* mice. At 3 weeks post injection, activation of hM3Dq by clozapine-N-oxide (CNO) was validated by testing activity-dependent cFos expression and action potential firing in Crabp1 neurons (Figures 3A, 3B and S4A). To assess the ability of Crabp1 neurons in driving energy expenditure, we deprived food from virally-transduced mice before administering CNO to rule out the effect of diet-induced thermogenesis (Figure 3C). Our results demonstrated that chemogenetic activation of Crabp1 neurons increased energy expenditure, oxygen consumption, and carbon dioxide release, independent of light or dark phases (Figures 3D, 3E, and S4B-S4D). We also found that administering CNO rapidly enhanced physical activity under both light and dark conditions (Figures 3F and 3G). In the control group of mice injected with AAV-DIO-mCherry, CNO treatment had no significant effect on energy expenditure or physical activity (Figures S4E and S4F). The rectal temperature and heat production by BAT were elevated by activating Crabp1 neurons in mice (Figures 3H and 3I). Consistent with BAT surface temperature increase, there was a prompt upregulation of *Ucp1* within BAT after neuronal activation (Figures 3J and S4G). Of interest, acute activation of Crabp1 neurons also increased feeding frequency without altering meal size and glucose metabolism (Figures 3K-3M and S4H-S4M). Therefore, we proposed a “mirror imbalanced model” wherein activating Crabp1 neurons enhances both energy expenditure and food intake, while deactivating — through disruption of synaptic transmission or chemogenetic inhibition — produces the opposite effect, albeit to potentially different extents (Figures 3N and S4N-S4R). In contrast, AgRP and POMC neurons constitute a functional unit to regulate energy intake and expenditure in an up-and-down manner, substantiating the “seesaw model”.

**Figure 3.**
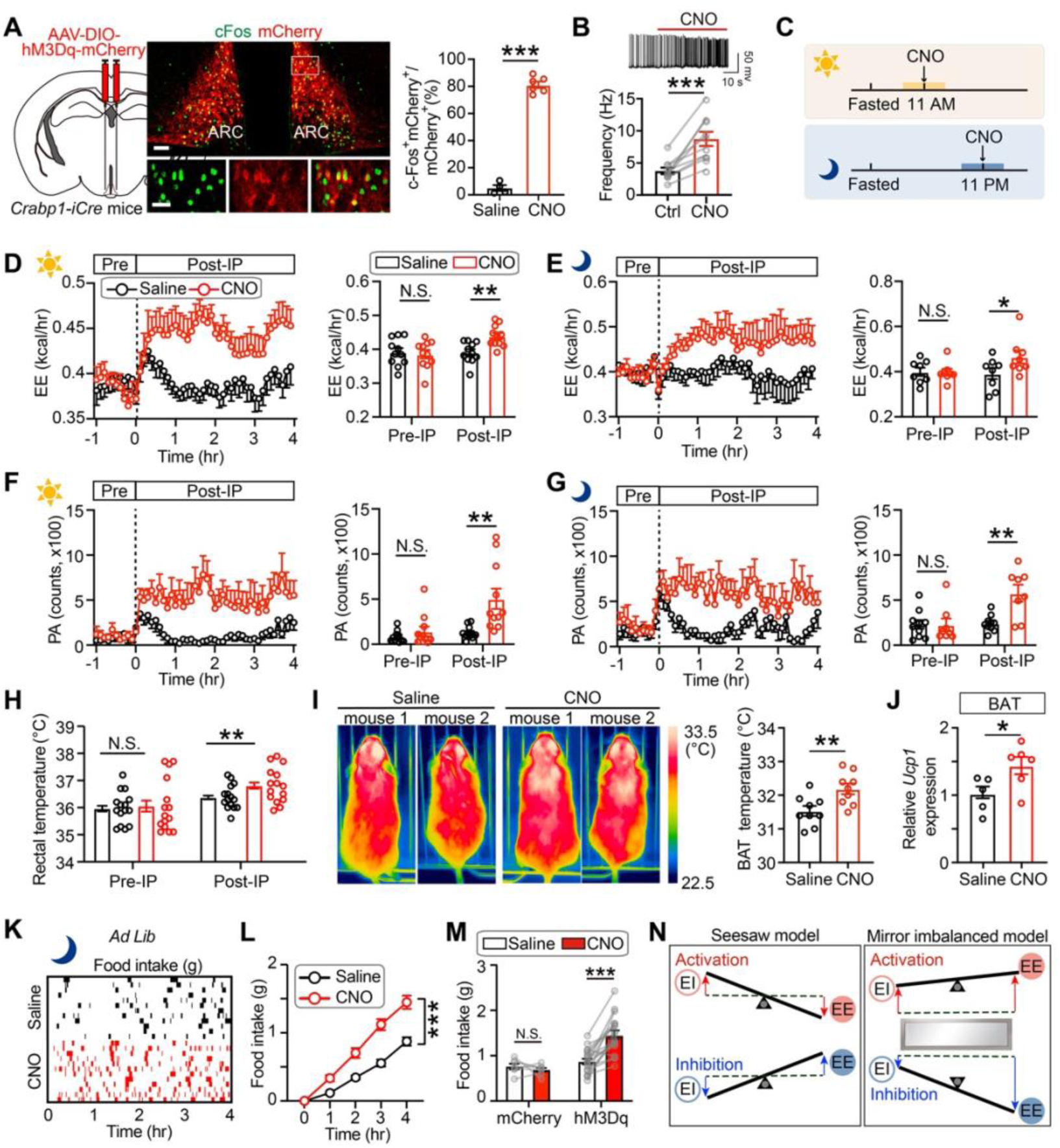
Chemogenetic activation of Crabp1 neurons promotes energy expenditure. (**A**) Histological validation of chemogenetic activation of Crabp1 neurons. Left: Schematic showing viral injection strategy for chemogenetic activation. Middle: Sample confocal images of cFos immunostaining among the mCherry^+^ Crabp1 neurons after clozapine-N-oxide (CNO) administration, with magnified views of the boxed regions shown below. Scale bars, 50 μm (top) and 20 μm (bottom). Right: Quantification of the proportion of cFos^+^ neurons among mCherry^+^ Crabp1 cells (n = 4 brains for saline group; n = 6 brains for CNO-activated group). (**B**) Electrophysiological validation of chemogenetic activation of Crabp1 neurons. Top: Sample electrophysiological trace showing the effect of CNO on the firing rate of Crabp1 neurons. Bottom: Quantitative analysis of action potential firing frequency in Crabp1 neurons before and after CNO perfusion on brain slices (n = 10 cells). (**C**) Experimental paradigm of chemogenetic manipulation of Crabp1 neurons during light and dark cycles. (**D** and **E**) Mean curves (left) and bar plots (right) showing real-time EE before (Pre-IP) and after (Post-IP) intraperitoneal injection of CNO (n = 8∼11) or saline (n = 8∼11) during light (D) and dark (E) phases. (**F** and **G**) Mean curves (left) and bar plots (right) showing real-time locomotion activity before and after intraperitoneal injection of CNO (n = 8∼11) or saline (n = 8∼11) during light (D) and dark (E) phases. (**H**) Quantification of rectal temperature before and after CNO or saline administration (n = 15). (**I**) Representative infrared thermograph images (left) and quantification of BAT temperature (right) at 4 hours post CNO (n = 9) or saline (n = 9) injection. (**J**) Gene expression analysis of *Ucp1* levels in BAT (n = 5 for saline group; n = 6 for CNO-activated group). (**K** and **L**) Raster plots (K) and mean curves (L) showing the dynamics of food consumption in *ad libitum*-fed mice within a 4 hours observation window following CNO or saline injection during the dark cycle. These mice expressed hM3Dq in Crabp1 neurons (n = 12∼15). (**M**) Quantification of 4 hours food intake after CNO or saline administration during the dark cycle. Control mice were microinjected with AAV-DIO-mCherry, while another group of *Crabp1-iCre* mice were injected with AAV-DIO-hM3Dq-mCherry (n = 7 for control group; n = 15 for hM3Dq group). (**N**) Schematics of the traditional “seesaw model”, exemplified by the role of AgRP neurons in EE and energy intake (EI), and our “mirror imbalanced model”. Values represent mean ± SEM. Significance was analyzed by paired or unpaired two-tailed Student’s *t* test and two-way ANOVA with Sidak’s multiple comparison test. N.S., not significant; *, *P* < 0.05; **, *P* < 0.01; ***, *P* < 0.001.

### Neural dynamics of Crabp1 neurons during cold exposure and physical activity

To decipher whether Crabp1 neurons modulate energy expenditure in response to external and internal challenges, we performed immunostaining to examine cFos expression following cold exposure, voluntary exercise, fasting, refeeding, and physical restraint (Figure S5A). Our quantitative analysis demonstrated that Crabp1 neurons were specifically activated during a 2 hours cold challenge at 4°C and 30 minutes of voluntary wheel running, with increased proportion and number of cFos-positive Crabp1 neurons compared to controls (Figures 4A and S5B); however, they did not respond to fasting, refeeding or restraint stress (Figures S5C and S5D). Further fluorescent ISH analysis revealed that high-fat diet (HFD) failed to change their response to fasting or refeeding (Figures S5E and S5F), suggesting that Crabp1 neurons may not sense or encode hunger and satiety.

**Figure 4.**
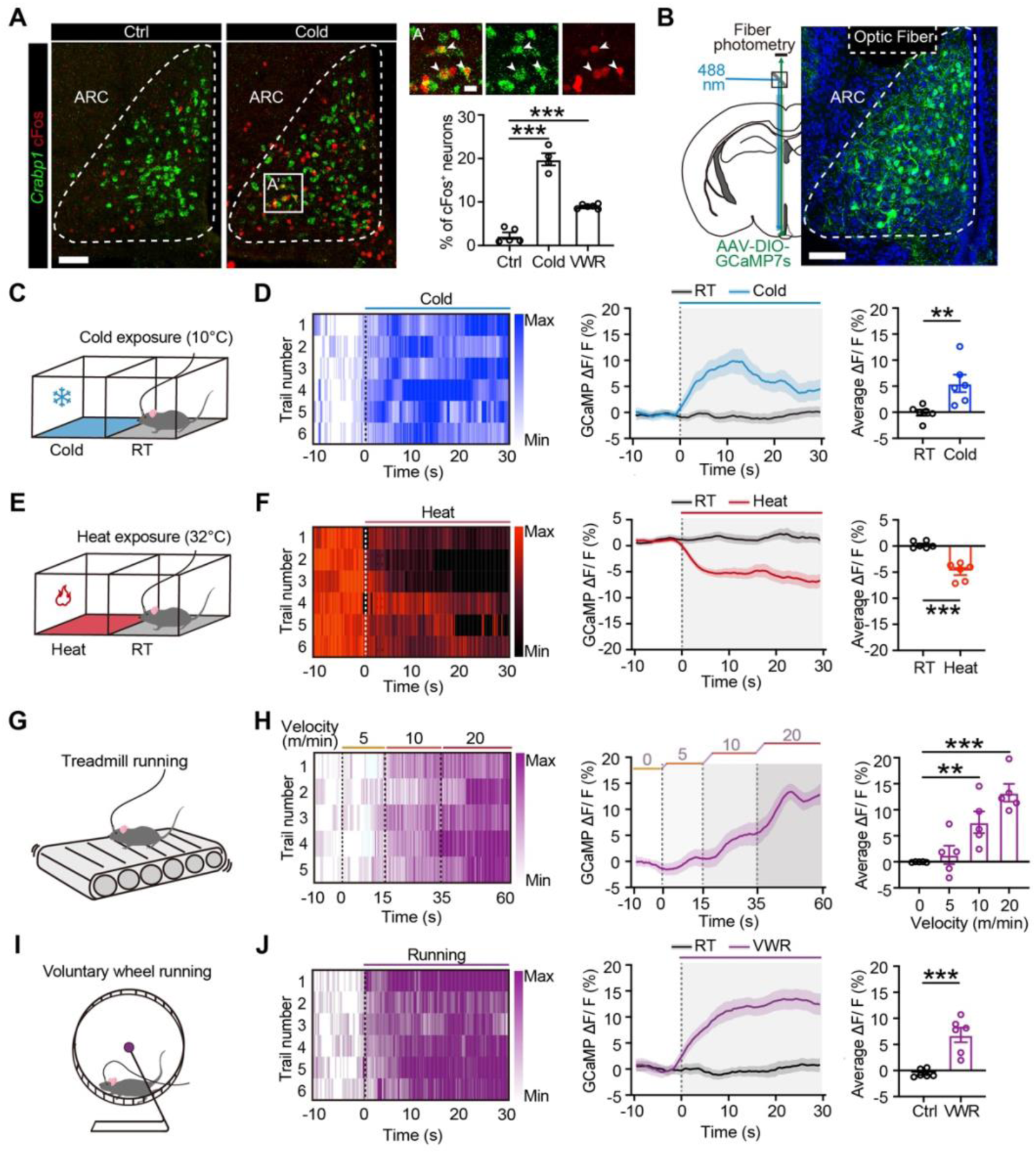
Crabp1 neurons respond to cold exposure and physical activity. (**A**) Activation of Crabp1 neurons by cold exposure and voluntary wheel running (VWR). Left: Representative confocal images showing the cFos immunoreactivity and *Crabp1* mRNA puncta in the ARC of control and cold-exposed mice. Scale bar, 50 μm. Top right: Magnified view of boxed area on the left showing the coexpression of cFos and *Crabp1* transcripts, indicated by white arrowheads. Scale bar, 10 μm. Bottom right: Quantification of the proportion of cFos^+^ neurons coexpressing *Crabp1* mRNA in the ARC of wild-type mice following cold exposure or VWR (n = 4∼6). (**B**) GCaMP7s expression in Crabp1 neurons. Left: Experimental strategy of viral injection of AAV-DIO-GCaMP7s and surgical implantation of optical cannula. Right: Sample image showing the expression of GCaMP7s in ARC Crabp1 neurons. Scale bar, 50 μm. (**C**) Schematic of the thermal shuttle box, with one chamber set to room temperature (RT) and the other chamber set to a cold environment at 10°C. (**D**) Neural dynamics of Crabp1 neurons during cold exposure. Left: Heatmap showing normalized calcium transient dynamics in Crabp1 neurons of individual mice in response to cold exposure. Middle: Mean calcium traces illustrating Crabp1 neuron response during RT and cold exposure. The change in GCaMP fluorescence (ΔF/F) reflects neuronal activity. Right: Box plots showing the average calcium response of Crabp1 neurons during RT and cold exposure (n = 6). (**E**) Schematic of another thermal shuttle box with one chamber set at 32°C. (**F**) Neural dynamics of Crabp1 neurons in a thermoneutral environment (n = 6). (**G**) Schematic of a mouse on the exercise treadmill. (**H**) Monotonic calcium response in Crabp1 neurons evoked by accelerating treadmill running. Left: Heatmap showing normalized calcium dynamics in individual mice during treadmill exercise with stepwise accelerating speed. Middle: Mean calcium traces illustrating Crabp1 neuron response during treadmill running. Right: Box plots showing the average calcium response of Crabp1 neurons at different treadmill running speed (n = 5). (**I**) Schematic of a mouse running on a voluntary wheel. (**J**) Calcium dynamics of Crabp1 neurons during VWR (n = 6). Values represent mean ± SEM. Significance was analyzed by paired two-tailed Student’s *t* test and one-way ANOVA with Dunnett’s multiple comparison test. **, *P* < 0.01; ***, *P* < 0.001.

For real-time recording the activity dynamics of Crabp1 neurons, we expressed GCaMP7s in these neurons via viral delivery and employed fiber photometry to measure their calcium transients (Figures 4B and S5G). Consistent with cFos staining results, neither access to food pellet after fasting nor ghrelin injection triggered calcium activity in Crabp1 neurons (Figures S5H and S5I). Given the induction of cFos in Crabp1 neurons after cold exposure, we connected two temperature-controlled chambers with a sealable corridor and found an increase in GCaMP fluorescence when mice entered the 10°C chamber from the 25°C chamber (Figures 4C and 4D). Conversely, GCaMP fluorescence signals in Crabp1 neurons declined as the mice shuttled from the 25°C chamber to a thermoneutral environment set at 32°C (Figures 4E and 4F).

As physical activity also contributes to energy expenditure, we measured calcium dynamics of Crabp1 neurons during both treadmill and voluntary wheel running. We found that physical exercise on treadmill evoked strong GCaMP response in Crabp1 neurons, which responded monotonically to stepwise increasing speed (Figures 4G and 4H). Similarly, Crabp1 neurons were activated by voluntary exercise on running wheels (Figures 4I and 4J). In the control group expressing EGFP in Crabp1 neurons, we observed no change in fluorescence during exposure to cold, heat, or treadmill running (Figures S5J and S5K). Together, these results suggest that Crabp1 neurons responded to external stimuli (e.g., cold exposure) and internal stimuli (e.g., physical exercise) to potentially increase energy supply and expenditure.

### Structural connectivity of Crabp1 neurons within and beyond the hypothalamus

To map the output connectivity of Crabp1 neurons, we first injected Cre-dependent AAV encoding membrane-tethered EGFP (mEGFP) into the ARC of *Crabp1-iCre* mice (Figures 5A and 5B). Structural analysis at 3 weeks post injection showed that Crabp1 neurons predominantly projected to two basal forebrain nuclei [i.e., ipsilateral lateral septum (LS) and bed nucleus of the stria terminalis (BNST)], as well as multiple hypothalamic nuclei [i.e., medial preoptic area (MPOA), paraventricular nucleus (PVN), lateral hypothalamus (LH), and dorsomedial nucleus (DMH)] (Figures 5C and 5D). To facilitate the visualization of axon terminals and synaptic boutons, we introduced AAV-FLEX-mGFP-2A-Synaptophysin-mRuby into Crabp1 neurons and similarly found a dense terminal labeling of LS, BNST, MPOA, PVN, LH, and DMH (Figures 5D and S6A-S6C).

**Figure 5.**
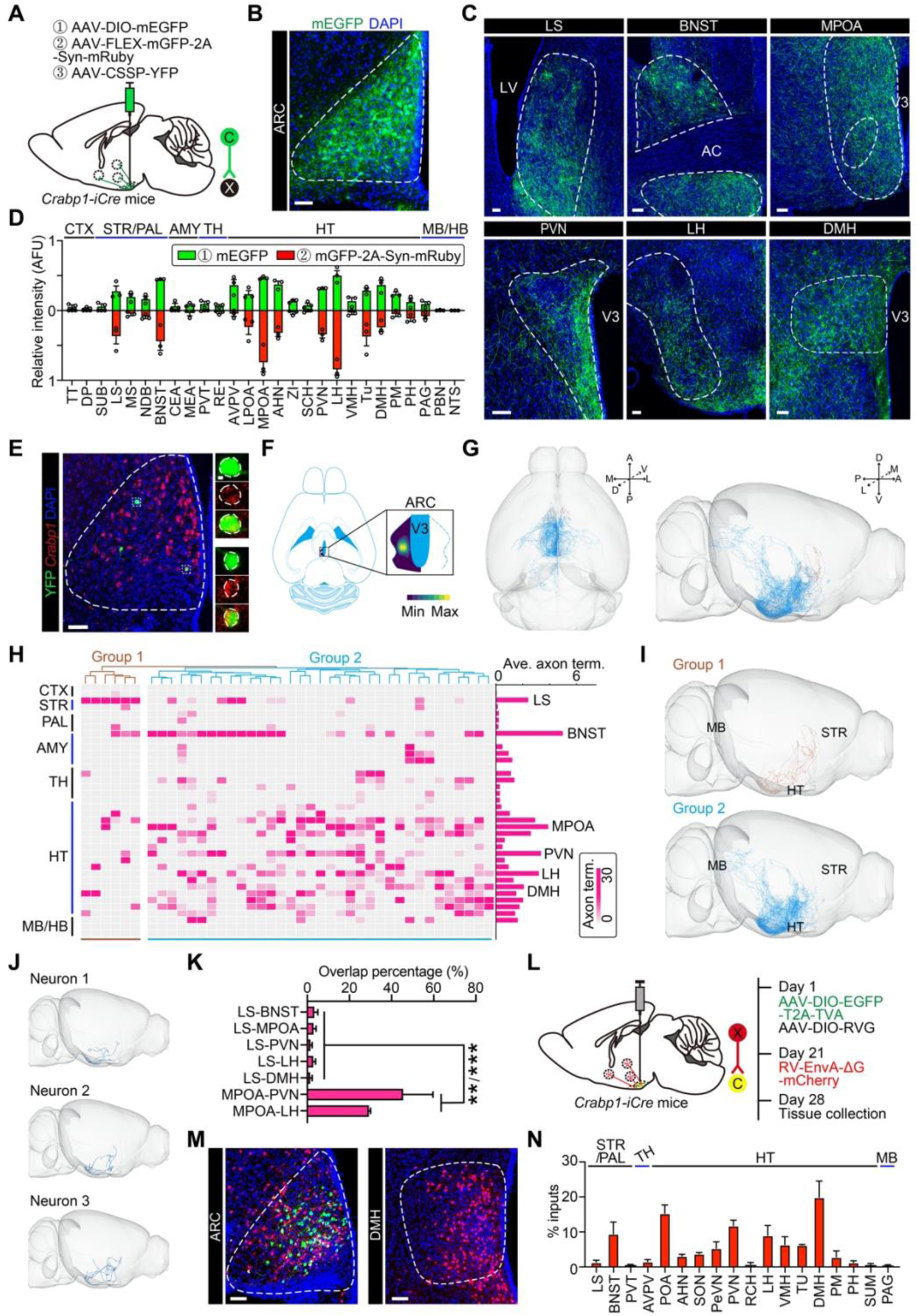
Mapping the input and output connectivity of Crabp1 neurons. (**A**) Schematic diagram showing three strategies of viral labeling of Crabp1 neurons for anterograde tracing: 1) AAV-DIO-mEGFP; 2) AAV-FLEX-mGFP-2A-Synaptophysin-mRuby; and 3) AAV-CSSP-YFP. FLEX, flip-excision switch; CSSP, cell type-specific sparse labeling. (**B**) Sample image showing the genetic labeling of Crabp1 neurons by mEGFP. Scale bar, 50 μm. (**C**) Representative confocal images showing the projection of mEGFP^+^ axonal fibers in multiple downstream target brain areas. See Table S6 for abbreviations. Scale bars, 50 μm. (**D**) Quantitative analyses of mEGFP^+^ axonal fibers (top) and mRuby^+^ axonal boutons (bottom) in various targeting brain regions. See Table S6 for abbreviations. Values represent mean ± SEM (n = 3 brains for each strategy). (**E**) Sample image showing the sparse labeling of Crabp1 neurons with YFP in the ARC, with magnified boxed regions highlighted on the right. *Crabp1* transcripts were detected by smFISH. Scale bars, 50 μm. (**F**) Exclusive distribution of Crabp1 neuronal soma within the ARC. Left: horizontal view of adult mouse brains. Right: Heatmap depicting the spatial distribution of labeled Crabp1 neurons for axonal tracing. (**G**) Three-dimensionally reconstructed neurons registered to the standard Allen Mouse Brain Atlas are shown in horizontal (left) and sagittal (right) views. A, anterior; P, posterior; D, dorsal; V, ventral; M, medial; L, lateral. (**H**) Heatmap demonstrating the regional distribution of axonal terminals for individual Crabp1 neurons. Hierarchical analysis identifies two groups. (**I**) Three-dimensional view of two Crabp1 neuronal groups with different axonal projection patterns. (**J**) Sample images showing the morphology of three reconstructed group 1 neurons. (**K**) Quantification of the proportion of overlapping neurons projecting to different downstream targets. (**L**) Experimental paradigm for monosynaptic retrograde tracing using a combination of AAV helpers and pseudotyped, glycoprotein-deleted rabies virus (RV-EnvA-ΔG). (**M**) Representative images showing the dually-labeled starter neurons in the ARC and mCherry^+^ upstream neurons in the DMH. Scale bars, 50 μm. (**N**) Quantification of the proportion of mCherry^+^ neurons in various brain regions, which monosynaptically project to the Crabp1 neurons, relative to all retrogradely-labeled cells. Values represent mean ± SEM (n = 3 brains). Significance was analyzed by one-way ANOVA with Dunnett’s multiple comparison test. **, *P* < 0.01; ***, *P* < 0.001.

The proposed one-to-one projection pattern of both AgRP and POMC neurons, which comprise distinct subpopulations projecting to separate downstream targets,^31,32^ prompted us to investigate the circuit organization of Crabp1 neurons at single-cell level. We administered AAV-CSSP-YFP to sparsely label Crabp1 neurons and imaged their fine morphology for tracing axonal projection based on fluorescence micro-optical sectioning tomography (fMOST; Figures 5E, 5F and S6D). After acquiring whole-brain images, we employed a standardized data processing workflow for brain-wide neuronal reconstruction and registered all traced projections onto the Allen Mouse Brain common coordinate frame (Figure 5G). While the majority of Crabp1 neurons innervated multiple targets via axon collateral branches, clustering analysis identified two distinct groups (Figure 5H). Group 2 neurons exhibited a more robust intra-hypothalamic projection and extended their axons beyond the hypothalamus into the BNST, whereas Group 1 neurons densely innervated into the LS with minimal projection to hypothalamic nuclei (Figures 5I and 5J). To cross-validate the anatomical organization of Crabp1 neurons, we injected retrograde AAVs carrying Cre-dependent EGFP and tdTomato into two distinct downstream target regions, and then examined the overlap between Crabp1 neurons projecting to these areas. Consistently, there was minimal overlap between LS-projecting Crabp1 neurons and those projecting to the MPOA, LH, PVN, DMH, or BNST, while we observed a robust co-labeling between MPOA-projecting neurons and those targeting the PVN and LH (Figures 5K and S7A-S7E). The divergent axonal projection patterns implicate a functional difference between these two groups of neurons and support a “one-to-many” wiring configuration (Figure S7F).

Furthermore, we compared the axonal projection pattern of Crabp1 neurons with AgRP and POMC neurons to illustrate their morphological differences. At the population level, the innervation of central amygdala, paraventricular thalamus, and parabrachial nucleus by Crabp1 neurons was barely detectable (Figures 5D, S6B and S6C), contrasting with the prominent projection of AgRP neurons to these brain regions (Figures S8A and S8B). As per the single-cell projectomic dataset, our quantitative analysis demonstrated that Crabp1 neurons displayed fewer axonal branches and terminals as well as shorter axon length and lesser innervation into the extrahypothalamic nuclei than AgRP and POMC neurons (Figures S8C-S8E). Thus, our data shed light on the predominant intrahypothalamic connectivity of Crabp1 neurons.

To identify the circuitry upstream of Crabp1 neurons, we determined to perform monosynaptic retrograde tracing with EnvA-pseudotyped, glycoprotein-deleted rabies virus (RV-EnvA-ΔG). Recombinant AAV expressing Cre-dependent avian-specific retroviral receptor and rabies virus glycoprotein were delivered into the ARC of *Crabp1-iCre* mice, followed by microinjection of RV-EnvA-ΔG-mCherry (Figure 5L). After 7 days, we mapped the upstream input neurons labeled with mCherry across the entire brain (Figure 5M). Systematic quantification showed multiple deep brain nuclei that were directly connected to Crabp1 neurons, including the DMH, POA, PVN, BNST, and LH (Figure 5N). Taken together with the output connectivity, our findings suggest that Crabp1 neurons establish reciprocal connections with neurons located within the hypothalamus and adjacent basal forebrain regions. Interestingly, the DMH and POA serve as critical components of the neural network controlling thermoregulation,^33,34^ while neuronal subsets in the PVN and LH have been shown to respond to physical activity and promote locomotion.^35–37^ Altogether, our results support the hypothesis that Crabp1 neurons may integrate temperature- and locomotion-related information and provide closed-loop feedback signals to control energy expenditure.

### Functional circuits of Crabp1 neurons in energy expenditure control

The intricate network established by Crabp1 neurons, which innervate multiple targets via collateral branches, suggests that a single group of Crabp1 neurons might regulate all facets of energy expenditure. Alternatively, it is plausible that distinct groups of Crabp1 neurons project to different downstream targets for controlling various components of energy expenditure. To test the hypothesis, we selected six brain regions with the highest density of axonal projections from Crabp1 neurons for functional validation, including the LS, BNST, MPOA, PVN, LH, and DMH. We simultaneously delivered retrograde AAV encoding Cre-dependent Flpo recombinase (AAV-Retro- FLEX-Flpo) to one of the downstream projection targets and AAV expressing Flpo- dependent TeNT (AAV-fDIO-TeNT-mCherry) to the ARC of *Crabp1-iCre* mice (Figure 6A). This strategy enabled us to preferentially inactivate neurons that project to specific downstream targets. At 4 weeks post viral delivery, we observed weight gain in mice following synaptic silencing of Crabp1 neuron subsets with biased innervation toward the BNST, MPOA, PVN, LH or DMH, with the exception of LS (Figure 6B). Consistent with the increase in the body weight and fat mass, we observed that synaptic inactivation of the axon terminals in the LH, MPOA, PVN, DMH, and BNST elicited reductions in total energy expenditure, oxygen consumption, physical activity, and cold-induced thermogenesis (Figures 6C-6F, S9 and S10). Genetic silencing any one of these five neuronal subsets was not sufficient to change the rectal temperature (Figures 6G, S9N, S9Y, S10K and S10V), implicating that shifting the core temperature set- point requires a large pool of Crabp1 neurons. The absence of change in body weight, fat storage, and energy expenditure components upon inactivating LS-projecting Crabp1 neurons supported the functional distinction between Group 1 and Group 2 neurons (Figures 6H-6L and S10W-S10Y). Summing up the phenotypic consequences of manipulating synaptic activity in the axon terminals led us to conclude that BNST-, MPOA-, PVN-, LH-, and DMH- but not LS-projecting Crabp1 neurons coordinate to regulate various aspects of energy expenditure (Figure 6M).

**Figure 6.**
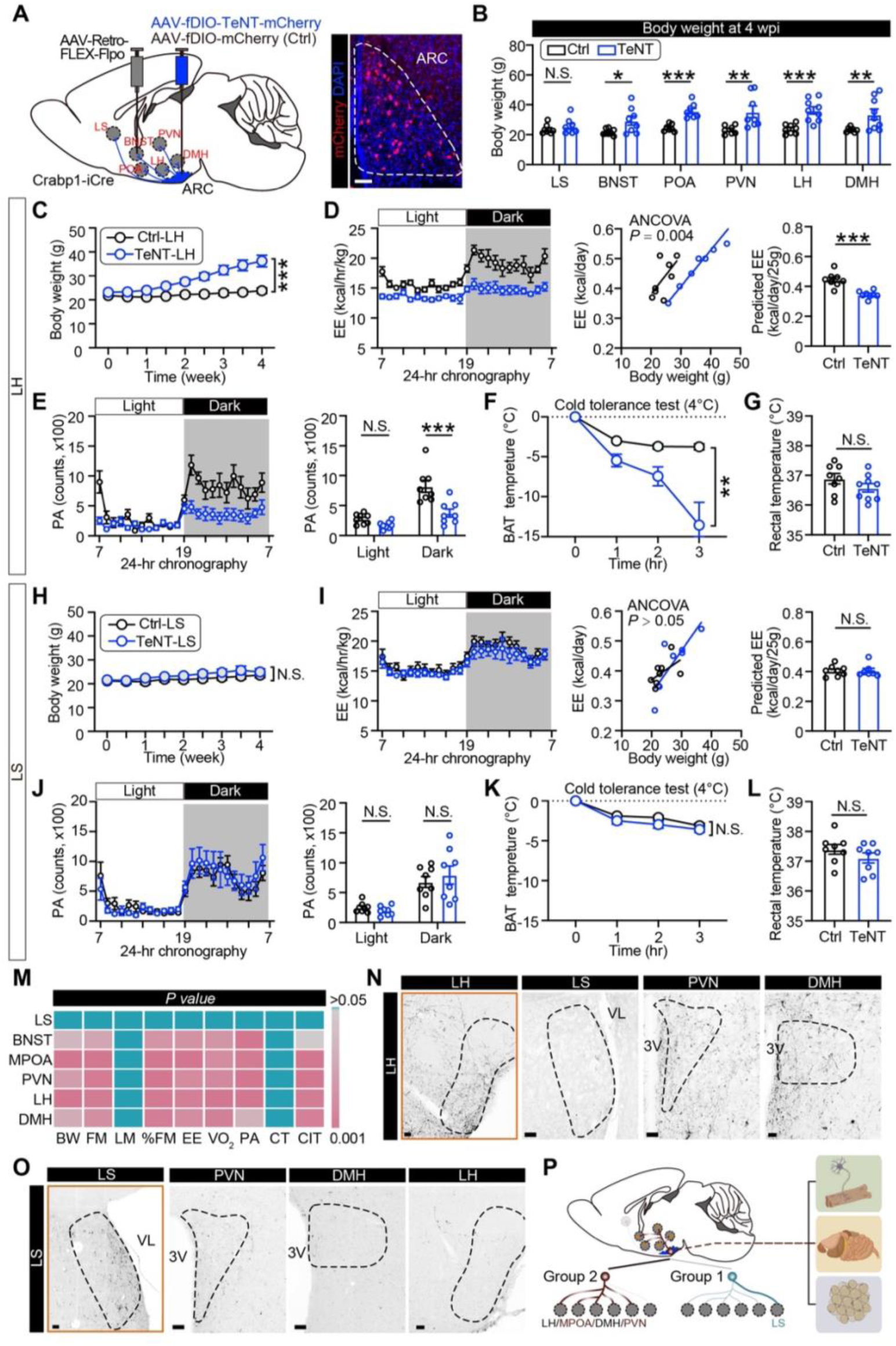
Crabp1 neurons coordinate energy expenditure via axon collateral branches. (**A**) Strategy for selectively silencing Crabp1 neuron subsets projecting to LS, BNST, MPOA, PVN, LH, or DMH. Left: Schematic diagram illustrating the delivery of retrograde AAV expressing Cre-dependent Flpo to one of the six projection targets and AAV encoding Flpo-dependent TeNT-mCherry to the ARC of *Crabp1-iCre* mice. Right: Sample image showing the labeling of a subset of Crabp1 neurons by mCherry in the ARC. Scale bar, 50 μm. (**B**) Quantification of body weight in control and TeNT-perturbed mice at 4 weeks post viral injection (wpi). Subsets of Crabp1 neurons projecting to LS, BNST, MPOA, PVN, LH, or DMH were synaptically inactivated by selective TeNT expression, with mCherry expression used as controls (n = 7∼9). (**C**-**G**) Body weight changes (C), total EE (D), locomotion activity (E), adaptive BAT thermogenesis during cold exposure (F), and rectal temperature (G) in mice subjected to synaptic silencing of the ARC^Crabp1^-LH pathway (n = 8∼9). Animals receiving AAV-fDIO-mCherry injection in the ARC served as controls. (**H**-**L**) Body weight changes (H), total EE (I), locomotion activity (J), rectal temperature (K), and adaptive BAT thermogenesis during cold exposure (L) in mice subjected to synaptic silencing of the ARC^Crabp1^-LS pathway (n = 8∼9). (**M**) Summary of the statistical analyses of metabolic phenotype following synaptic inactivation of Crabp1 neuron subsets targeting LS, BNST, MPOA, PVN, LH, or DMH (n = 8∼9). BW, body weight; FM, fat mass; LM, lean mass; % FM, percentage of fat mass; VO2, oxygen consumption. (**N** and **O**) Representative images showing the distribution of mCherry^+^ axonal fibers upon injection of AAV-fDIO-mCherry into the ARC of *Crabp1-iCre* mice and AAV-Retro-FLEX-Flpo into LH (N) or LS (O), two of Crabp1 neuron efferent sites (denoted with orange boxes). Scale bars, 50 μm. (**P**) Schematic model summarizing that Group 2 Crabp1 neurons, rather than Group 1, coordinate multiple components of EE. Values are presented as mean ± SEM. Statistical significance was analyzed by two-way ANOVA with Sidak’s multiple comparison test or unpaired two-tailed Student’s *t* test. Statistical analysis in panels D and I were performed using regression-based ANCOVA with EE as dependent variable and body mass as covariate. N.S., not significant; *, *P* < 0.05; **, *P* < 0.01; ***, *P* < 0.001.

To further characterize the axonal collateralization of these neuronal subsets, we analyzed the distribution of mCherry^+^ axon terminals throughout the entire brain. Consistent with the neuronal tracing data (Figure 5H), we observed that neuronal subsets projecting toward the LH, MPOA, PVN, and DMH extended collateral branches to one another (Figures 6N and S11). In contrast, LS-targeting Crabp1 neurons displayed a much less extensive intra-hypothalamic projection (Figure 6O). In combination with the structural projections of Crabp1 neurons, these results strongly suggest that Group 2 neurons, rather than Group 1, play a pivotal role in coordinating energy expenditure and controlling body weight (Figure 6P).

### Prolonged light exposure compromises Crabp1 neuron-mediated energy expenditure

The role of Crabp1 neurons in energy consumption raises the possibility that their disrupted neuronal activity in specific environmental contexts may contribute to metabolic disease progression. Prolonged light exposure, a consequence of the widespread adoption of artificial light in modern society, has been linked to the global increase in the prevalence of human metabolic disorders such as obesity and diabetes.^38,39^ Rodents exposed to extended daylight periods exhibited attenuated energy expenditure and increased adiposity without altering food intake.^40,41^ Importantly, photoperiodic regulation of the hypothalamic *Crabp1* gene expression in seasonal animals implies that Crabp1 neuronal activity may be susceptible to prolonged light exposure.^42,43^ Toward this end, we subjected adult wild-type mice to two different daily light exposure regiments for 5 weeks (Figure 7A). Mice in the long day (LD) group with 18 hours daily light gained more body weight and fat mass compared to those in the equal day (ED) with 12 hours daily light (Figures 7B and 7C). Our calorimetry analysis revealed that daily energy expenditure was declined after a period of prolonged light exposure (Figure 7D), as supported by previous findings.^40,41^ To determine the potential effect of long photoperiod on Crabp1 neurons, we microinjected AAV-DIO-mCherry into the ARC of *Crabp1-iCre* mice, exposed the mice to LD or ED, and subsequently performed perforated patch-clamp recordings (Figure 7E). Importantly, we found that the average firing rate of spontaneously active Crabp1 neurons was reduced following prolonged light exposure (Figures 7F and 7G). The proportion of spontaneously silent neurons was also increased by the long photoperiod (LD, 51.6%; ED, 22.5%), and the membrane potential was hyperpolarized (Figures 7H and 7I). These results strongly suggest that increased daylight exposure disrupted the spontaneous firing of Crabp1 neurons.

**Figure 7.**
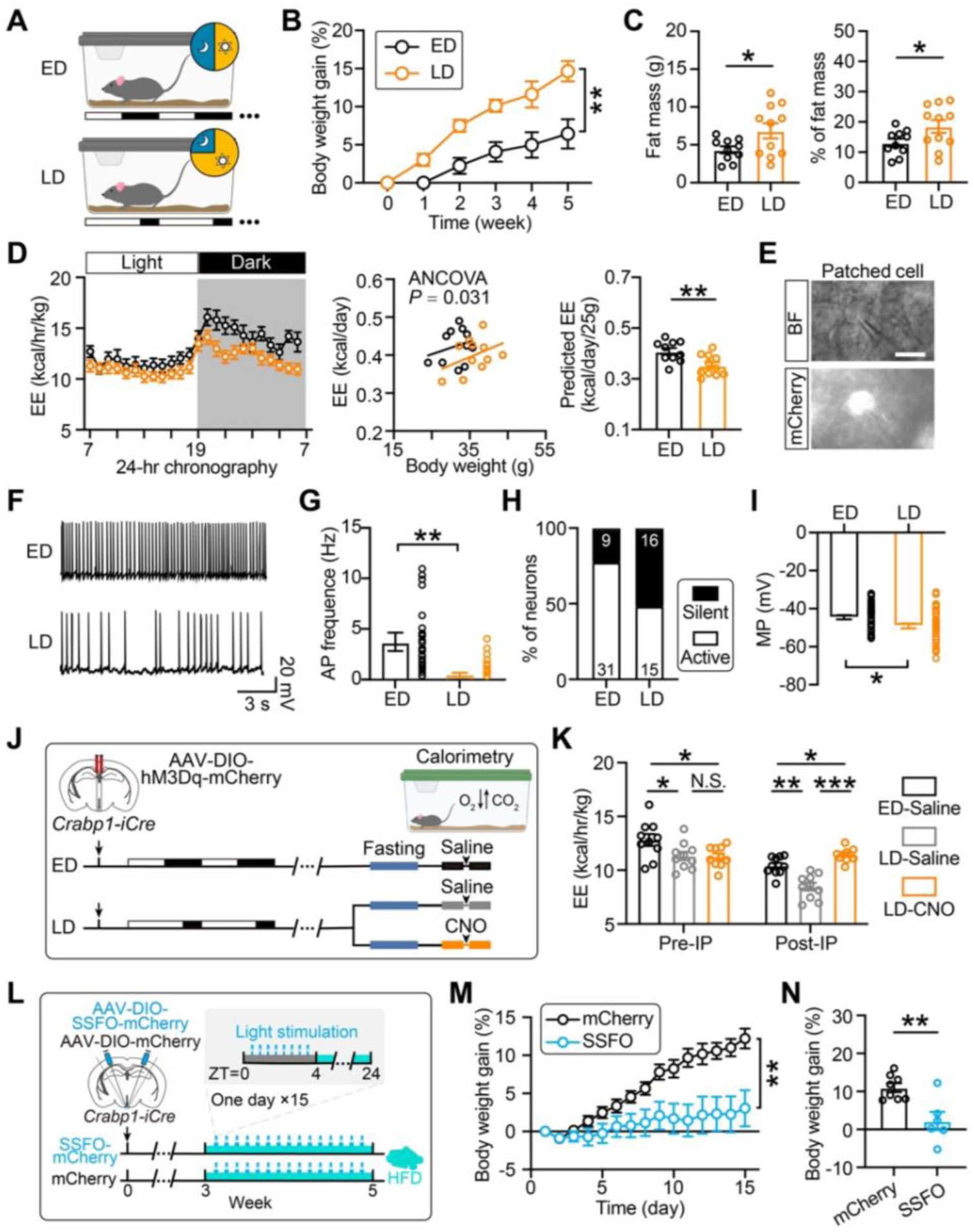
Susceptibility of Crabp1 neurons to long photoperiods and their activation in mitigating diet-induced obesity. (**A**) Schematic illustrating the strategy for comparing the metabolic profiles of mice exposed to either 12 hours or 18 hours daily light cycles. ED, equal day (12-hour light:12-hour dark); LD, long day (18-hour light: 6-hour dark). (**B**) Body weight gain observed over a 5-week period with ED or LD (n = 10 for ED group; n = 11 for LD group). (**C**) Quantification of fat mass (left) and its proportion relative to body weight (right) in mice exposed to ED or LD (n = 10∼11). (**D**) Total EE (left), EE regression against body weight (middle), and predicted EE with 25g body weight (right) for mice housed in different daily photoperiods (n = 10∼11). Statistical analysis was performed using regression-based analysis of ANCOVA with EE as dependent variable and body mass as covariate. (**E**) Sample image showing the expression of mCherry in a patched cell. BF, bright field. Scale bar, 10 μm. (**F**) Representative electrophysiological traces showing the spontaneous action potential firing pattern of the Crabp1 neurons in mice with ED and LD exposure. (**G**) Quantification of action potential firing frequencies in Crabp1 neurons from mice exposed to different durations of daily light (n = 40 for ED group; n = 31 for LD group). (**H**) Proportion of spontaneously active and silent (≤ 0 Hz) Crabp1 neurons. Empty and filled bars denote active and silent cells, respectively. Absolute numbers of neurons are indicated. (**I**) Measurement of membrane potential in Crabp1 neurons from mice in ED (n= 40) and LD (n= 31) groups. (**J**) Schematic illustrating the experimental design for analyzing the effect of Crabp1 neuron activation on the EE of mice exposed to long photoperiods. Animals receiving CNO injection in the ED group and saline injection in the LD group were used as controls. (**K**) Quantitative analysis of 4 hours EE before and after administration of CNO or saline in mice from the ED and LD groups (n = 10∼11). (**L**) Schematic showing the experimental design for chronic optogenetic activation of Crabp1 neurons. The mice were fed with HFD at 3 weeks after viral transduction and subjected to 4 hours of daily blue light activation, with food deprivation during the optogenetic stimulation. (**M**) Body weight changes in control (n = 8) and optogenetically-manipulated (n = 6) mice over a 15-day period. (**N**) Quantification of body weight change in control (n = 8) and optogenetically-activated (n = 6) mice at 15 days post-treatment. Values are presented as mean ± SEM. Statistical significance was analyzed by two-way ANOVA with Sidak’s multiple comparison test, one-way ANOVA with Dunnett’s multiple comparison test, and paired or unpaired two-tailed Student’s *t* test. N.S., not significant; *, *P* < 0.05; **, *P* < 0.01; ***, *P* < 0.001.

To assess whether chemogenetic activation of Crabp1 neurons could rescue the prolonged light-induced decline in energy expenditure, we microinjected AAV-DIO-hM3Dq-mCherry into the ARC of *Crabp1-iCre* mice and randomly assigned them into either ED or LD exposure group. After 5 weeks, we conducted calorimetry analysis and collected data before and after intraperitoneal CNO injection (Figure 7J). As anticipated, LD-treated mice displayed lower energy expenditure compared to the ED group prior to intraperitoneal injection. However, CNO administration successfully reversed this decrease in energy expenditure in LD mice, whereas saline injection had no effect (Figure 7K). Thus, these data collectively suggest that prolonged light exposure compromises Crabp1 neuronal activity and thereby reduces the total energy expenditure, which may increase the risk of metabolic disorders.

### Chronic activation of Crabp1 neurons prevents high-fat diet-induced weight gain

Given the role of Crabp1 neurons in energy expenditure, we next sought to investigate whether their chronic activation could sustain elevated energy expenditure and counteract weight gain in pathological conditions, such as the widely used high-fat diet-induced obesity model. To achieve sustained activation of Crabp1 neurons, we employed stabilized step-function opsins (SSFOs) for optogenetic manipulation, which enables long-lasting neuronal activation with brief light stimulation while eliminating the need for repeated CNO injections required for chemogenetics.^44^ Following bilateral viral transduction of SSFO into the ARC of *Crabp1-iCre* mice and optical fiber implantation, the mice were fed with HFD to induce obesity and subjected to 4 hours of daily blue light activation of Crabp1 neurons at 3 weeks post-injection (Figure 7L). Over a 15-day treatment period, we found that optogenetic stimulation of Crabp1 neurons significantly mitigates HFD-induced body weight gain, as compared to the control mice (Figures 7M and 7N). These results imply that Crabp1 neurons may serve as a potential target for preventing HFD-induced obesity.

## Discussion

Maintenance of energy homeostasis and body weight requires a balance between energy intake and expenditure. Disruption of energy homeostasis due to excessive calorie intake and/or reduced energy expenditure has contributed to the global obesity crisis.^45,46^ While initial weight loss has been achieved by restricting caloric intake (e.g., dieting and GLP1 receptor agonist),^47,48^ rebound weight gain remains a major issue due to synaptic adaptations of the hunger circuits as a counter-regulatory mechanism.^21^ To circumvent this challenge, understanding the mechanisms of balanced energy expenditure is critical for developing long-lasting therapeutic interventions. However, the neural basis for regulating energy expenditure in the brain remains poorly understood. In this study, we explored the molecular, structural, and functional characteristics of a poorly understood population of “non-AgRP, non-POMC” neurons - specified by *Crabp1* expression. We reveal that arcuate Crabp1 neurons and their extensive intra-hypothalamic projections via collateral branches are critical for energy expenditure, physical activity, and thermoregulation. The mode of Crabp1 neurons regulating energy balance conform to “mirror imbalanced model”, rather than the traditional “seesaw model”. We further demonstrate the photoperiod-dependent susceptibility of Crabp1 neurons and their potential role in ameliorating HFD-induced obesity.

The ARC serves as a critical node of hypothalamic control of energy balance.^49,50^ Arcuate neurons, primarily categorized as glutamatergic and GABAergic, can be iteratively refined into additional subtypes. For example, glutamatergic neurons include POMC and KNDy neurons, with POMC neurons further being split into three distinct subtypes.^11^ Here, we revealed that arcuate GABAergic neurons comprised of *Agrp*-, *Sst*-, *Ghrh*-, *Th*- and *Crabp1*-expressing subtypes. Dual-color *in situ* hybridization demonstrated that Crabp1 neurons show little with POMC, AgRP, KNDy, SST, GHRH and *Th*^+^*Slc6a3*^+^ dopaminergic subtypes, supported by single-cell profiling of the entire hypothalamus.^23,51^ Functional enrichment analysis unveiled that genes specific to Crabp1 neurons are linked to cell adhesion, retinoic acid metabolism, thyroid hormone signaling, and neurotransmitter receptor activity, implying robust cell-cell communication within the Crabp1 neuronal network. In contrast to AgRP and POMC neurons, Crabp1 neurons exhibited low levels of *Lepr* and *Insr* expression, as supported by previous *in situ* hybridization results.^24^ Instead, *Thrb* was robustly expressed in this neuronal subtype, suggesting its potential modulation by systemic or local thyroid hormones. Overall, we established a systematic categorization of arcuate neurons, uncovering a functionally-undefined neuronal subtype and elucidating its molecular characteristics.

At the functional level, hypothalamic AgRP and POMC neurons act in an opposing manner in perceiving systemic energy status and operate in a “seesaw model” to regulate energy balance.^3,52^ Activation of POMC neurons contributes to a stable body weight by suppressing food intake and enhancing energy dissipation, while activation of AgRP neurons stimulates food intake and reduces energy expenditure during the hunger state.^53,54^ However, this conventional perspective may still be incomplete and overly simplistic. Here, we demonstrated that inactivation of Crabp1 neurons reduces energy expenditure, while chemogenetic activation displays the opposite effect. Based on the finding that synaptic silencing of Crabp1 neurons leads to the onset of obesity, we propose a “mirror imbalanced model” wherein the extent to which Crabp1 neurons regulate energy expenditure and food intake may vary. During winter or hunting, animals likely require increased energy expenditure to engage in activities such as foraging or combating cold temperature.^55,56^ Thus, Crabp1 neurons may have evolved in response to these challenges to increase energy expenditure. Similar to most arcuate GABAergic neuronal subtypes, such as AgRP, SST, TH and PNOC neurons,^7,11,57,58^ Crabp1 neuronal activation also stimulates food intake. The abundance of neuronal subtypes promoting food intake may have evolved as redundant mechanisms in response to periods of food scarcity. Unlike other arcuate GABAergic subtypes, Crabp1 neuronal activation is not triggered by hunger, but is observed to regulate food intake, perhaps to preemptively compensate for future caloric deficit. Together, we hypothesize that distinct arcuate neuronal subpopulations regulate energy expenditure and intake in divergent and redundant ways, enabling animals to cope with various physiological conditions and environmental challenges. While neurons that conform to the “seesaw model” maintain energy homeostasis by toggling between states of hunger and satiety, those following the “mirror imbalanced model” may adapt to disadvantageous conditions to regulate the deployment of energy supply to enhance survival.

Neural circuit connections lay the structural foundation for exerting diverse physiological functions. Our single-neuron tracing data show that POMC and AgRP neurons display robust intra-hypothalamic connections as well as project toward central amygdala, paraventricular thalamus, periaqueductal gray and parabrachial nucleus, supporting previous studies using anterograde tracing at the population level.^59^ Although Crabp1 neurons exhibit similar intra-hypothalamic axonal projections as POMC and AgRP neurons, they display less complexity in axon terminal arborization and extra-hypothalamic projections. Of note, it has been suggested that the functional circuits of AgRP neurons are organized into a “one-to-one” wiring configuration, in which neurons are subdivided into distinct subsets with separate projections to individual downstream brain regions.^31,60^ For example, AgRP neurons stimulate food intake by targeting PVN, promote food seeking by projecting to paraventricular thalamus, inhibit territorial aggression by extending toward amygdala, and modulate pain through connections to parabrachial nucleus.^7,61–63^ In this study, we found Group 2 Crabp1 neurons may primarily promote overall energy metabolism via a “one-to- many” projection pattern where neurons are wired with multiple target regions with collateral branches. Multiple downstream targets of Crabp1 neurons, including PVN, DMH, LH, and MPOA, have been well-established to directly or indirectly regulate the autonomic nervous system to modulate energy expenditure, physical activity, and thermogenesis.^64,65^ While our mapping of single-neuron projection identifies two group of Crabp1 neurons, the specific function of Group 1 Crabp1 neurons remains to be explored.

Internal homeostasis of an organism is maintained through proactive adaptations of hypothalamic neurons to internal and external fluctuations.^66^ Hypothalamic Crabp1 neurons may perceive and integrate relevant stimuli for improved survival. For example, both cold exposure and physical activity stimulate Crabp1 neurons to drive energy expenditure to facilitate thermogenesis for cold defense and enhance exercise motivation. In contrast, AgRP neurons predominantly contribute to appetite control, but are not responsible for promoting thermogenesis or meeting energy demands under various stimuli.^67,68^ Moreover, we found that the activity of Crabp1 neurons is inhibited in thermoneutral environment and by prolonged light exposure; both conditions are known to increase body weight.^41,69^ The abrogation of Crabp1 neuron activity by long photoperiod exposure reduces energy expenditure and reactivation of Crabp1 neurons in these animals is sufficient to restore the light-induced reduction in energy expenditure. Given the association between light pollution and increased prevalence of metabolic disorders in humans, our study may provide a mechanistic insight into how nocturnal artificial light exposure increases the risk of obesity.^39,70^

Together, our study identifies Crabp1 neurons as a neural basis for regulating energy expenditure and propose a “mirror-imbalance model” in which hypothalamic neurons integrate environmental stimuli for coping with adverse conditions. The elucidation of their molecular and circuit profiles may open an alternative avenue for preventing metabolic disorders.

### Limitations of the study

There are certain limitations to our study. First, while previous studies have reported photoperiodic regulation of hypothalamic *Crabp1* gene expression in seasonal animals,^42,43^ we did not specifically examine the role of *Crabp1* in energy homeostasis. Second, our study did not explore the impact of sex on energy metabolism in Crabp1 neuron function.

## RESOURCE AVAILABILITY

### Lead Contact

Further information and requests for resources and reagents should be directed to and will be fulfilled by the Lead Contact, Qing-Feng Wu.

### Materials Availability

All animals and unique/stable reagents generated in this study are available from the Lead Contact with a completed Materials Transfer Agreement.

### Data and code availability

Single-nucleus RNA-seq data is accessible in National Center for Biotechnology Information (NCBI) under accession code: GSE276414. All data and computational code used in this work are available upon request. Any additional information required to reanalyze the data reported in this paper is available from the lead contact upon request.

## ACKNOWLEDGMENTS

We gratefully acknowledge Dr. Shengjin Xu and Rong Gong for comments on the manuscript. We would like to thank Dr. Kun Li for technical help. We thank bioRENDER (biorender.com) for assisting in drawing schematic diagrams. The work was supported by National Natural Science Foundation of China (32230031, 31921002, 32070972 and 81891002), National Key R&D Program of China (2019YFA0800213), Beijing Municipal Science & Technology Commission (Z210010), and Hundred-Talent Program (Chinese Academy of Sciences).

## AUTHOR CONTRIBUTIONS

Q.W. and T.W. designed all experiments. T.W. and S.H. performed all experiments as well as statistical analyses. Y.W., Z.C., and T.W. conducted bioinformatics analysis. X.L., Y.L., X.X., and H.G. produced and analyzed the single-neuron tracing data. S.Z., F.Z., M.J., and P.C. contributed to the electrophysiological recording. Q.L. provided the optogenetic virus. Z.X., J.W., and M.X. provided experimental assistance and technical advice. Q.W., T.W., and Y.Z.L. wrote the manuscript.

## SUPPLEMENTAL INFORMATION

**Document S1. Figures S1–S11**

**Table S1. Signature genes for each cell class, related to Figure 1**.

**Table S2. Signature genes for each neuronal subtype, related to Figure 1 and S2.**

**Table S3. Genes enriched in Crabp1 neurons, related to Figure 1 and S2.**

**Table S4. DNA sequences of *in situ* hybridization probes, related to Figure 1 and S1.**

**Table S5. DNA sequences of real-time quantitative PCR primers, related to Figure 2 and S3.**

**Table S6. Abbreviations of anatomic structures, related to Figure 2**.

## STAR METHODS

### KEY RESOURCE TABLE

#### EXPERIMENTAL MODEL AND SUBJECT DETAILS

##### Animals

*Crabp1-iCre* mouse strain (Stock No. T037951) was ordered from the Gempharmatech Co. Ltd. In brief, the sgRNA directs the Cas9 endonuclease to cleave the downstream sequence of the transcription start site of the *Crabp1* gene, resulting in the insertion of the iCre-stop element by homologous recombination. The inserted element is approximately 1.9 kb in length, and the iCre is expressed under the control of the endogenous *Crabp1* gene, followed by transcriptional termination of the *Crabp1* gene. *AgRP-IRES-Cre* (Stock No. 012899) and *POMC-Cre* (Stock No. 005965) mouse strains were obtained from the Jackson Laboratory. Wild-type C57BL/6N mice were ordered from SPF Biotechnology Co. Ltd (Beijing, China). Animals were maintained on a 12-hour light/12-hour dark cycle at a temperature of 22-25°C with 50-60% humidity and provided with food and water *ad libitum*. Male mice were used in all experiments unless otherwise stated. All experimental procedures were performed according to protocols approved by the Institutional Animal Care and Use Committee at the Institute of Genetics and Developmental Biology, Chinese Academy of Sciences.

In a long photoperiod experiment, mice from the same litter aged 8-10 weeks were randomly assigned to two groups: equal day (ED, 12-hour light/12-hour dark cycle) and long day (LD, 18-hour light/6-hour dark cycle), for a treatment period of 5 weeks. During the acclimation period, mice had *ad libitum* access to standard laboratory chow and water. All cabinets were kept in the same room, sharing an air supply, and maintained at the same temperature, humidity, and light intensity (∼ 20 μW/cm^2^). The photoperiods were adjusted so that lights turned on simultaneously for both the ED and LD groups, promoting synchronization at the start of the daily circadian rhythm.

## METHOD DETAILS

### Single nucleus RNA library preparation and sequencing

The hypothalamic arcuate nucleus (ARC) was microdissected from adult mice under a stereoscope, immersed in ice-cold homogenization buffer containing 250 mM sucrose, 25 mM KCl, 5 mM MgCl_2_, 10 mM Tris-HCl, 0.1 mM dithiothreitol (DTT), 1% bovine serum albumin (BSA), 0.1% NP-40, and ribonuclease inhibitor (0.4 U/μL), and triturated using a glass homogenizer for 10 times. After centrifugation of homogenized tissues at 500 g for 5 minutes, we washed the cell pellets with homogenization buffer using repetitive pipetting and isolated the nuclei through a subsequent centrifugation step. The resulting cell pellets were suspended in cold phosphate-buffered saline (1×PBS) to generate single-nucleus suspensions for single-nucleus RNA-seq (snRNA-seq). snRNA-seq libraries were prepared utilizing DNBSEQ technology platforms (BGI Genomics, China) and the DNBelab C4 Single-Cell Library Prep Set (MGI Tech, #1000021082). In brief, single-nucleus suspensions were pumped through a microfluidic device to generate droplets. After breaking the emulsion and collecting the beads, reverse transcription and cDNA amplification were carried out to produce barcoded libraries. The resulting cDNA was then fragmented into short pieces measuring 250 to 400 bp, and indexed sequencing libraries were constructed following the manufacturer’s protocol. The quality of the sequencing libraries was evaluated using the Qubit ssDNA Assay Kit (Thermo Fisher Scientific, #Q10212). The barcoded libraries were loaded into patterned nanoarrays and sequenced using the ultrahigh-throughput DIPSEQ T-series sequencer.

### Sequencing data preprocessing

The raw sequencing reads from the DIPSEQ T-series sequencer were filtered and demultiplexed using PISA (version 0.10; https://github.com/shiquan/PISA). Reads were aligned to GRCm38 mouse genome using STAR (version 2.7.4a) and sorted by Sambamba (version 0.7.0). PISA was utilized to generate a nucleus versus gene unique molecular identifier (UMI) count matrix for downstream analysis.

#### Ambient RNA removal

Ambient RNA noise was mitigated using SoupX (version 1.4.8; https://github.com/constantAmateur/SoupX), with default settings except for the contamination fraction (denoted as rho). The rho value was automatically parameterized using the autoEstCont function in tissues where rho was lower than 0.05 or higher than 0.2, and manually set to 0.2 using the setContaminationFraction function if the autoEstCont value was between 0.05 and 0.2.

#### Quality control and data integration

For quality assurance, cells were screened based on the following criteria prior to further analysis: (i) the percentage of mitochondrial gene counts was required to be less than 10%; (ii) nuclei needed to have detected gene counts between 200 and 4,000; and (iii) RNA counts had to be less than 10,000 to meet the quality criteria. The matrix files arising from different batches of experiments were subsequently integrated and analyzed using Seurat software (version 4.0.1).

#### Dimensionality reduction and clustering

After removing unqualified cells, we normalized the raw reads using the “LogNormalize” subroutine with 10,000 scales, selected the top 2,000 variable genes using the “FindVariableFeatures” function, and rescaled the normalized dataset with the “ScaleData” function in Seurat software. Principal components analysis was performed to reduce the dimensionality of the datasets, and the first 30 principal components were further analyzed using t-distributed stochastic neighbor embedding (t-SNE) for dimensionality reduction. We further performed clustering analysis using a shared nearest neighbor (SNN) based clustering algorithm, which calculates *K*-nearest neighbors (*K* = 25), constructs an SNN graph, and optimizes the modularity function to determine clusters. For cell clustering, we adopted stepwise clustering strategy for each dataset. First, the cells passing quality control were clustered with the Louvain algorithm in “FindClusters” function at a resolution of 0.5 to distinguish neurons from nonneuronal cells. Subsequently, we subsetted neurons from diverse single-cell datasets for further reclustering. The neuronal clusters were further compared pairwise to identify cell type-specific genes, and cell identities were assigned by cross-referencing their marker genes with known neuronal subtype markers.

To analyze the molecular features of Crabp1 neurons, we compared their transcriptome with other arcuate neurons and identified differentially expressed genes using the "FindMarkers" function in Seurat with the recommended settings (Table S2). Moreover, we performed an enrichment analysis of molecular features of Crabp1 neurons using the "GSEA" (gene set enrichment analysis) function in clusterProfiler with hallmark gene sets from the Molecular Signatures Database (Table S3) and visualized the results using the enrichplot package. In addition, the protein-protein interaction network of Crabp1 neuronal molecular features was constructed using STRING (https://string-db.org/), followed by *k*-means clustering (N = 5) to reveal distinct functional clusters within the network. The pseudobulk expression analysis was conducted using Seurat’s "AverageExpression" function, providing insights into aggregated gene expression patterns across cellular populations.

#### Single-molecule fluorescent in situ hybridization

To collect adult mouse brains, we anesthetized mice by intraperitoneal injection of 2,2,2-tribromoethanol (Sigma, #T48402; 400 mg/kg), and performed transcardial perfusion with saline followed by 4% paraformaldehyde (PFA) in 1×PBS. Mouse brains were dissected, post-fixed for 8 hours in 4% PFA at 4℃, and subsequently cryo- protected in 20% sucrose in 1×PBS for 12 hours followed by 30% sucrose for 24 hours. Tissue blocks were prepared by embedding in Tissue-Tek O.C.T. Compound (Sakura, #4583). The brain sections (20-40 mm in thickness) were prepared using a cryostat microtome (Leica, CM3050S) and stored in -80℃ freezer.

For single-molecule fluorescent *in situ* hybridization (smFISH) by RNAscope technology, the probes targeting against *Crabp1* (#474711-C3), *Slc32a1* (#319191-C2) and *Fos* (#316921-C1) were designed by Advanced Cell Diagnostics. Briefly, the brain sections were dried at 50℃ for 2 hours, rinsed with 1×PBS, treated with 3% hydrogen peroxide in methanol and underwent antigen retrieval. Subsequently, tissue sections were dehydrated with 100% ethanol and incubated with mRNA probes for 2 hours at 40℃. Specific signals were then amplified with multiplexed amplification buffer according to the manufacturer’s protocol (Advanced Cell Diagnostics, #323110) and detected using TSA Plus Fluorescence kits (PerkinElmer, #NEL753001KT).

For smFISH by hybridization chain reaction (HCR) approach, we designed HCR probes by MATLAB codes (https://github.com/GradinaruLab/HCRprobe) for targeting the mRNA sequence of *Crabp1*, *Agrp*, *Pomc*, *Th*, *Ghrh*, *Tac2*, *Sst*, *Slc17a6*, *Gad1* and *Gad2* genes. All HCR probes were synthesized by Sangon Biotech and dissolved in DEPC-H_2_O (Table S4). Brain sections were permeabilized in 70% ethanol for 12-16 hours at 4°C, followed by 0.5% Triton X-100 in 1×PBS at 37°C for 1 hour, and treated with protease K (Invitrogen, #AM2546; 10 μg/mL) to improve mRNA accessibility. After two washes with 1×PBS at room temperature (RT), sections were prehybridized in 30% probe hybridization buffer for 10 minutes at 37°C and then incubated in 30% probe hybridization buffer containing HCR probes (10 mM for each) at 37°C for 3-5 hours. After mRNA hybridization, the washing and amplification steps were performed as previously described.^71^ Immunofluorescence staining with antibodies against Fos was further conducted on HCR-labeled sections.

#### Immunohistochemistry

For immunostaining, the tissue sections were washed with 1×PBS, then antigen retrieval was performed at 95°C by 1×target retrieval solution (DAKO, #S1699), and pre-blocked with 1×TBS++ (TBS containing 5% donkey serum and 0.3% Triton X- 100) for 1 hour at RT, followed by incubation with primary antibodies diluted in TBS++ overnight at 4°C. After primary antibody incubation, the brain sections were washed for 3 times with 1×PBS and incubated with the secondary antibodies for 2 hours at RT. The slides were mounted with medium containing 120 mg/mL polyvinyl alcohol and 31.25 mg/mL anti-fluorescence quencher 1, 2-dianilinoethane. The primary antibodies used included rabbit anti-Fos (Synaptic systems; #226008; 1:500), goat anti-GFP (Rockland; #600-101-215; 1:1,000), and rabbit anti-RFP (Rockland; #600-401-379; 1:1,000). The secondary antibodies used were anti-goat/rabbit Cy2, anti-goat/rabbit Cy3, and anti-goat/rabbit Cy5 (Donkey; Jackson ImmunoResearch; 1:500).

#### Viruses

The following viral tools were used in this study: AAV2/9-hEF1a-DIO-mCherry-P2A- TetTox-WPRE-pA (1.02×10^13^, Taitool, S0506-9); AAV2/9-hSyn-DIO-mCherry-WPRE-pA (1.93×10^13^, Taitool, S0240-9); AAV2/9-hSyn-DIO-hM3Dq-mCherry- WPRE-pA (1.04×10^13^, Taitool, S0192-9); AAV2/9-hSyn-DIO-hM4Di-mCherry- WPRE-pA (1.25×10^13^, Taitool, S0193-9); AAV2/9-hSyn-DIO-jGCaMP7s-WPRE-pA (1.23×10^13^, Taitool, S0590-9); AAV2/9-hSyn-DIO-EGFP-WPRE-pA (1.78×10^13^, Taitool, S0746-9); AAV2/9-hSyn-FLEX-mGFP-2A-Synaptophysin-mRuby-WPRE-pA (1.25×10^13^, Taitool, S0250-9); AAV2/9-hSyn-DIO-mEGFP-WPRE-pA (2.78×10^12^,BrainVTA, PT-3478); AAV2/2Retro-hSyn-FLEX-EGFP-WPRE-pA (1.93×10^13^, Taitool, S0239-2R); AAV2/2Retro-hSyn-FLEX-tdTomato-WPRE-pA (1.56×10^13^, Taitool, S0255-2R); AAV2/9-EF1a-DIO-RVG-WPRE-hGH-pA (1.53×10^12^, BrainVTA, PT-0023); AAV2/9-EF1a-DIO-EGFP-T2A-TVA-WPRE-hGH-pA (1.53×10^12^, BrainVTA, PT-0062); RV-ENVA-ΔG-mCherry (2.00×10^8^,BrainVTA, R01004); AAV2/2Retro-CAG-FLEX-Flpo-WPRE-pA (2.08×10^13^, Taitool, S0273-2R) and AAV2/9-hEF1a-fDIO-mCherry-P2A-TetTox-WPRE-pA (1.34×10^13^, Taitool, S0551-9). To achieve specific sparse labeling of neuronal projections, we employed AAV2/9-CSSP-YFP-8E3 (5.43×10^12^, Brain Case, BC-SL003), which was produced by co-packaging the AAV2/9-CMV-DIO-Flpo plasmid and AAV2/9-fDIO-EYFP plasmid with the ratio of 1:8000 in a single rAAV production step. AAV-PHP.eB-EF1a-DIO-SSFO-mCherry was provided by Qinghua Liu at National Institute of Biological Sciences. All viruses were aliquoted and stored at -80℃ before use.

#### Stereotaxic injection

7-12 weeks old mice were anesthetized with 1.5-3% isoflurane and placed on a stereotaxic frame (RWD). Body temperature was maintained at 35-37°C by a heating instrument (Reptizoo). Erythromycin eye ointment was applied to the eyes, which were then covered with tin foil. Virus was injected with a volume of 100-200 nL/site at a rate of 25 nL/min using a micropump system (KDS, Legato 130) with a glass capillary. When the infusion was done, the glass pipette was kept inserted for 10 minutes before being slowly withdrawn to prevent backflow of fluids. To recover from anesthesia, the mice were placed in a clean cage on a heating pad until they regained consciousness. Mice were kept in the cage for at least 2 weeks for recovery and viral expression before behavioral testing.

For neural manipulation, 200 nL AAV2/9-hSyn-DIO-hM3Dq-mCherry-WPRE-pA or AAV2/9-hSyn-DIO-hM4Di-mCherry-WPRE-pA were injected bilaterally into the ARC of *Crabp1-iCre* mice. Antero-Posterior (AP) -1.45 mm, mediolateral (ML) ±0.25 mm, dorsoventral (DV) -5.85 mm. Control group was injected with mCherry virus (AAV2/9-hSyn-DIO-mCherry-WPRE-pA). For chronic synaptic silencing, 200 nL AAV2/9-hEF1a-DIO-mCherry-P2A-TetTox-WPRE-pA was injected as described above. We also bilaterally injected 200 nL AAV-PHP.eB-EF1a-DIO-SSFO-mCherry or AAV2/9-hSyn-DIO-mCherry-WPRE-pA into the ARC at a 10-degree angle for optogenetic stimulation. Two optic fibers (diameter, 200 μm; NA, 0.37; Inper) were then implanted at the same angle, 0.2 mm above each injection site.

For fiber photometry recording, 200 nL AAV2/9-hSyn-DIO-jGCaMP7s-WPRE-pA or AAV2/9-hSyn-DIO-EGFP-WPRE-pA were injected unilaterally into ARC of *Crabp1-iCre* mice. And then an optic fiber (diameter, 200 μm; NA, 0.37; RWD) was implanted to the same coordinate as that of the virus injection. The fiberoptic cannula was affixed to the skull with dental universal light curing microglass composite (Charisma).

For the axon terminal labeling, 300 nL AAV2/9-hSyn-DIO-mEGFP-WPRE-hGH- pA and AAV2/9-hSyn-FLEX-mGFP-2A-Synaptophysin-mRuby-WPRE-pA were injected unilaterally into the ARC of *Crabp1-iCre* mice. Mice were kept in the cage for at least 4.5 weeks for viral expression prior to sacrifice for anterograde tracing. We further microinjected 200 nL AAV2/2Retro-hSyn-FLEX-EGFP-WPRE-pA and AAV2/2Retro-hSyn-FLEX-tdTomato-WPRE-pA into the LS, BNST, MPOA, PVN, LH, and DMH of *Crabp1-iCre* mice for retrograde tracing. Mice were sacrificed after 3 weeks to allow for viral expression.

For neural circuit inhibition, 200 nL AAV2/2Retro-CAG-FLEX-Flpo-WPRE-pA was injected bilaterally into lateral septum (LS; AP 0.5 mm, ML ±0.5 mm, DV -2.75 mm) and 200 nL of AAV2/9-hEF1a-fDIO-mCherry-P2A-TetTox-WPRE-pA was injected into ARC of *Crabp1-iCre* mice. The control group was injected with AAV2/9- hEF1a-fDIO-mCherry-WPRE-pA. For the different nuclei, the coordinates were respectively bed nucleus of the stria terminalis (BNST; AP 0.05 mm, ML ±0.85 mm, DV -3.65 mm), medial preoptic area (MPOA; AP 0.14 mm, ML ±0.3 mm, DV -4.9 mm), paraventricular nucleus (PVN; AP -0.9 mm, ML ±0.25 mm, DV -4.8 mm), lateral hypothalamus (LH; AP -0.94 mm, ML ±0.7 mm, DV -5.1 mm), dorsomedial nucleus (DMH; AP -1.7 mm, ML ±0.4 mm, DV -4.9 mm).

For rabies-mediated monosynaptic retrograde tracing, 200 nL of virus mixture containing AAV2/9-EF1a-DIO-RVG-WPRE-hGH-pA and AAV2/9-EF1a-DIO-EGFP-T2A-TVA-WPRE-hGH-pA was injected unilaterally into ARC of *Crabp1-iCre* mice. Three weeks later, 200 nL of RV-ENVA-ΔG-mCherry was injected into the ARC at the same site and the mice were sacrificed for histological analyses a week later.

#### Fiber photometry recording

Fiber photometry experiments were started at least 3 weeks after AAV2/9-hSyn-DIO- jGCaMP7s-WPRE-pA or AAV2/9-hSyn-DIO-EGFP-WPRE-pA injection. The implanted optic fiber was previously connected to a fiber photometry system (ThinkerTech, China) via an optical fiber patch cord. To record fluorescence signals, a beam from a 480 LED was reflected with a dichroic mirror, focused with a lens coupled to a complementary metal oxide semiconductor (CMOS) detector. The LED power ranged from 0.03 to 0.05 mW at the tip of the optical fiber patch cord to minimize bleaching. The mice must be acclimated to the chamber and connected with optical fiber patch cord for more than 5 minutes before testing different cues.

To cold or hot stimuli, mice were placed in a custom-made two-room acrylic chamber (50×25×25 cm^3^). One side of the chamber has a room-temperature floor plate and the other side has either a 10℃ or 32℃ floor plate. Mice were first acclimated in the room-temperature chamber for 5 minutes, afterwards, the door between the chambers was removed to allow the mice to explore the cold or hot test chambers. Once the mice entered the test room, the door in the middle was shut to confine the mice in the test chamber for no less than 2 minutes. The first step of the mice into the test room was taken as the onset of the event.

For treadmill running test, the mice were trained to learn the treadmill three days prior to the experiment. On the test day, mice were placed on the treadmill and acclimated to it for the first 5 minutes, following which the mice were subjected to the treadmill running. Each exercise section consisted of three stages: (stage 1) a 15 seconds run at a low speed of 5 m/min, (stage 2) another 15 seconds at a medium speed of 10 m/min, and (stage 3) another 15 seconds at a high speed of 20 m/min in which the treadmill speed gradually increased from 5 to 10 m/min or from 10 to 20 m/min over 5 seconds.

For voluntary wheel running, the mice were habituated to a rotating wheel in their home cages for at least a week before the experiment. On the test day, mice were placed in their home cages with a rotating wheel for 30 min. Voluntary running was recorded using a video camera and manually analyzed to extract the time points for movement onsets.

For food pellet test, the mice were food-deprived for 24 hours. On the test day, the mice were placed in home cages and a small food pellet (TestDiet) was left in the home cages every 5 minutes. The food pellet weighted only 45 mg to ensure that the mice could eat it in one bite, with the first bite serving as the event timestamp marker. Wood chips of the same size were used as control objects instead of the food pellets. The experiments were repeated 7-10 times per mouse.

For the ghrelin test, the mice were injected with saline as a control. The mice received a 1 mg/kg intraperitoneal injection of ghrelin (Tocris, #1465) and the moment they were returned to home cages marked the onset of the event. The mice were recorded for at least 5 minutes prior to the saline or ghrelin injection.

#### Metabolic phenotype analysis

For energy expenditure and physical activity measurement, mice were moved to an indirect calorimetry system (PhenoMaster, TSE system, Germany). Mice were acclimatized to the metabolic cages for at least 3 days prior to data collection. During the monitoring period, the mice were individually housed and had *ad libitum* access to food and water, with a 12-hour light/12-hour dark cycle and an ambient temperature of 22°C. After calibrating the system with the reference gases (20.95% O_2_ and 0.05% CO_2_), the oxygen consumption, carbon dioxide production, respiratory exchange ratio (RER = VCO_2_/VO_2_), energy expenditure, and locomotion activity were recorded. Mice injected with AAV2/9-hEF1a-DIO-mCherry-P2A-TetTox, AAV2/9-hEF1a-fDIO-mCherry-P2A-TetTox, and control virus were collected in the TSE system for three days. Mice injected with AAV2/9-hSyn-DIO-hM3Dq-mCherry, AAV2/9-hSyn-DIO- hM4Di-mCherry or AAV2/9-hSyn-DIO -mCherry were collected in the TSE system 1 hour pre-injection and 4 hours post-injection of saline or clozapine-N-oxide (CNO, Enzo Life Sciences, BML-NS105-0025). The ten-day experimental procedure began with three days of acclimatization to the metabolic cages and intraperitoneal saline injections to habituate the mice to handling and injections. After an overnight fast on the fourth day, a subset of mice was randomly assigned to receive either saline or CNO injections on the fifth day, with food access restored four hours later. A recovery period was allowed from days six to eight, followed by a second overnight fast on the ninth day. On the tenth day, the saline and CNO injections were reversed from Day 4, administered during either the light or dark cycle, and food access was resumed four hours after injection.

For body weight and composition measurement, mice were housed in groups (3-5 animals per cage) and fed a standard chow diet, with body weight recorded twice a week. Body composition, including fat mass and lean mass, of all mice was measured using a magnetic-resonance imaging analyzer (EchoMRI, USA). In optogenetic experiments, the mice were virally transduced with either AAV-DIO-SSFO-mCherry or AAV-DIO-mCherry and subjected to 4 hours of daily blue light activation (1 s 1 Hz, 1 s on /30 min off, 8-10 mW) over 15 consecutive days. Food was withdrawn during the light activation. Body weight was measured daily throughout the treatment period.

For food intake test, the mice were housed individually at least 7 days before the food intake assays. To prevent the accumulation of food debris, automatic feeders (BiolinkOptics) with small food pellets (TestDiet, 45 mg) were used for food intake calculation. In TeNT-induced synaptic silencing experiments, food intake was measured daily for 3 days. In chemogenetic experiments, food intake was measured after intraperitoneal injections of saline or CNO (1 mg/kg) for 4 hours.

#### Temperature Measurements

For core temperature measurement, rectal temperature was recorded using a thermometer (Kew Basis, FT3400). The thermometer probe, coated with petroleum jelly, was inserted into the anal canal to record the stabilized temperature hourly before and during exposure to 4°C. Excrement and urine of mice were cleaned to minimize experimental error. For chemogenetic experiments, rectal temperature was recorded 4 hours after CNO injection.

For brown adipose tissue (BAT) temperature measurement, an infrared thermograph camera (Fortric 225S) was used to detect the surface temperature above BAT from a top-down view. Each time the camera was positioned at the same distance from the mice to ensure that a clear image of the mouse body outline and hair could be captured. The heatmaps were automatically generated by the camera software. BAT surface temperature was measured hourly before and during exposure to 4°C, as well as 4 hours after CNO injection.

#### Glucose and insulin tolerance test

Prior to glucose tolerance test (GTT), mice underwent 16 hours fasting period, followed by an intraperitoneal injection of glucose at a dose of 2 g/kg. Blood glucose concentrations were measured at 5 min before and 15, 30, 60 and 120 min after glucose injection using Accu-Chek Active Blood Glucose Meter (Roche, Swiss). For the insulin tolerance test (ITT), mice fasted for 5 hours received an intraperitoneal injection of insulin (0.75 U/kg), and their insulin sensitivity was then assessed by measuring blood glucose levels as described above.

#### Hormonal assays

Animals were deeply anesthetized, and blood samples were allowed to clot before being centrifuged at 1,000 g for 20 min to obtain serum. The basal levels of mouse thyroid-stimulating hormone (TSH), tetraiodothyronine (T4), adrenocorticotropic hormone (ACTH) and cortisol (CORT) were quantified with enzyme-linked immunosorbent assay (ELISA) kits from Mlbio (#ml063200V, #ml001955V, #ml001895V, and #ml037564V). Briefly, we added 50 µL of serum samples and 100 µL of streptavidin-horseradish peroxidase conjugate into the designated wells of assay plates coated with specific antibodies, incubated the plates at 37°C for 1 hour on a plate shaker, and measured the absorbance of the reaction solution with a plate reader (SpectraMAX 190, Molecular Devices, USA). The hormone concentrations were determined using the standard curve.

#### Adipose tissue histology

Mice were deeply anesthetized with isoflurane, and BAT, inguinal white adipose tissue (iWAT), and epididymal white adipose tissue (eWAT) were dissected, fixed in 4% PFA overnight, embedded in paraffin, and sectioned at a thickness of 5 μm using a rotary microtome (Leica, RM2255). Histological changes were examined by hematoxylin and eosin (H&E, Beyotime, #C0105) staining and observed by Olympus VS200 Slide Scanner (Olympus Life Science, Japan). The size of adipose tissue was quantified using an automated plugin called Adiposoft in ImageJ.^72^

#### Quantitative real-time PCR

Real-time PCR (RT-qPCR) was performed using SYBR Green PCR Master Mix (Roche, #4913850001). Briefly, we extracted total RNA from tissues with Trizol reagent (Invitrogen, #15596018) and performed reverse transcription with M-MLV reverse transcriptase (Invitrogen, #28025013) to obtain cDNA. Real-time PCR reactions were carried out in triplicate in a reaction volume of 10 μL using 5 μL of 2×FastStart Universal SYBR Green Master mix, 0.5 μL of each primer, 1 μL of purified cDNA and 3 μL nucleic acid-free water. Cycling conditions consisted of an initial denaturation step of 95°C for 3 min followed by 40 cycles of 95°C denaturation for 10 seconds and an annealing/extension step of 30 seconds at 60°C. Changes in fold amplification were analyzed by the comparative ΔΔCt (cycle threshold) method. All primer sequences are listed in Table S5.

#### Fluorescence micro-optical sectioning tomography imaging and 3D visualization

For sparse labeling, we injected 100 nL of AAV-CSSP-YFP into the ARC of *Crabp1- iCre* mice. After 4.5 weeks, the mice were scarified, and virus-labeled brains were dehydrated with alcohol and embedded in resin. Whole-brain datasets were then acquired with the fluorescence micro-optical sectioning tomography imaging (fMOST) system. In short, the sample was fixed on the base, images of the top surface were captured using two fluorescent channels, and the imaged tissue was subsequently removed. This process allowed us to obtain continuous whole-brain datasets layer by layer with high resolution (0.32×0.32×1 μm^3^). To visualize the data in 3D and perform statistical analysis, we registered the datasets to the Allen Mouse Brain Common Coordinate Framework version 3 (CCFv3).^73^ Image preprocessing was utilized to correct uneven illumination and remove background noise. The downsampled data (with a voxel resolution of 10×10×10 μm^3^) was imported into Amira software (version 6.1.1) to distinguish and extract regional anatomical invariants, such as the brain outline, ventricles, and corpus striatum. Subsequently, a grey-level-based registration algorithm was applied to register the extracted features. Basic operations, including extraction of areas of interest, resampling, and maximum projection, were conducted using Amira software and ImageJ (version 2.1.0).

#### Morphological reconstruction of single neuron

For single-cell morphological analysis, we employed semi-automatic methods to reconstruct the morphology of sparsely labeled neurons, following previous studies.^74^ Initially, we obtained the spatial coordinates of labeled somas from high-resolution data and converted the data format of GFP-labeled data from TIFF to TDI type using Amira. Subsequently, the data block containing the designated soma was loaded into GTree software, where we assigned the soma as the initial point and marked all its fibers with unfinished tags. We then selected an uncompleted fiber and traced it in the next block using automatic tracing. After tracing, we reviewed the traced fiber and marked its branches with unfinished tags. This process was repeated until the selected fiber was completed, and then we reconstructed the remaining unfinished fibers until all fibers were completed. The reconstructed neurons were checked back-to-back by three persons. The tracing results were saved in SWC format. Additionally, we registered the propidium iodide labeled data and corresponding tracing results to the reference atlas.

#### Electrophysiological Recordings

Brain slices containing the ARC were prepared from adult mice under anesthesia induced by isoflurane before decapitation. The brains were swiftly extracted and immersed in ice-cold oxygenated cutting solution (228 mM sucrose, 5 mM glucose, 26 mM NaHCO_3_, 1 mM NaH_2_PO_4_, 2.5 mM KCl, 7 mM MgSO_4_, and 0.5 mM CaCl_2_) saturated with 95% O_2_ and 5% CO_2_. Using a vibratome (VT1200S, Leica), coronal brain slices (300 μm) were obtained. These slices were then incubated at 28°C in oxygenated artificial cerebrospinal fluid (ACSF: 119 mM NaCl, 2.5 mM KCl, 1 mM NaH_2_PO_4_, 1.3 mM MgSO_4_, 26 mM NaHCO_3_, 5 mM glucose, and 2.5 mM CaCl_2_) for 30 minutes, followed by an additional hour at room temperature under the same conditions before being transferred to the recording chamber. To reduce synaptic input, it contained 50 μM PTX (picrotoxin, P1675, Sigma Aldrich), 50 μM DL-AP5 (DL-2- amino-5-phosphonopentanoic acid, BN0086, Biotrend), and 20 μM CNQX (6-cyano- 7-nitroquinoxaline-2,3-dione, C127, Sigma-Aldrich). The ACSF was perfused at a rate of 1 mL/min. Visualization of the acute brain slices was achieved using a 40× Olympus water immersion lens, differential interference contrast optics (Olympus), and a charged-coupled device camera.

Patch pipettes were pulled from borosilicate glass capillary tubes (Warner Instruments, #64-0793) using a PC-10 pipette puller (Narishige). For recording of APs (current clamp), pipettes were filled with solution (135 mM K-methanesulfonate, 10 mM HEPES, 1 mM EGTA, 1 mM Na-GTP, 4 mM Mg-ATP and 2% neurobiotin, pH 7.4). The resistance of pipettes varied between 3.0 MΩ and 4 MΩ. The current and voltage signals were recorded with MultiClamp 700B and Clampex 10.5 data acquisition software (Molecular Devices). After establishment of the whole-cell configuration and equilibration of the intracellular pipette solution with the cytoplasm. Recordings with series resistances greater than 15 MΩ were rejected.

## QUANTIFICATION AND STATISTICAL ANALYSIS

### Fiber photometry data analysis

All data analyzed using MATLAB (R2022, MathWorks) codes. The fluorescence change (ΔF/F) was calculated as (F-F_0_)/F_0_, where F_0_ is the baseline fluorescence signal, following previous studies.^75^ The baseline was defined as the 10 seconds (e.g., cold exposure, heat exposure, treadmill running, voluntary wheel running or food pellets) or 1 minute (e.g., ghrelin injection) time window preceding the event onset.

#### Axonal fiber and synaptic boutons quantification

To quantify the density of Crabp1 and AgRP neuronal axon projections, we delineated the target areas in each region containing fibers according to the Allen Mouse Brain Atlas and calculated the average pixel intensity as F_raw_ using ImageJ. Additionally, a boxed area of the same size but in a brain region lacking fiber terminals was selected to calculate the background intensity (F_background_) on the same image. F_signal_ was subsequently calculated as F_raw_ - F_background_. For each animal, F_signal_ was normalized by the maximum F_signal_ in the ARC. Normalized F_signal_ was then used to calculate the average terminal field intensity across animals.

To quantify synaptic boutons of Crabp1 neurons in downstream brain regions, red (mRuby^+^ synaptic boutons) and blue (DAPI) channel TIFF images were imported into ImageJ. Maximum intensity projection along the Z-axis were generated for each channel, and exported as single-channel 16-bit gray scale high-resolution images. Structure boundary ROIs were delineated based on criteria from Allen Mouse Brain Atlas using the blue channel in QuPath (version 0.3.2). The identification and quantification of axonal boutons were conducted utilizing Otsu’s thresholding algorithm for segmenting and detecting individual signals, all achieved with CellProfiler (version 4.2.1). For quantitative analysis, every third section of each brain was selected.

#### Axonal terminals quantification and hierarchical clustering of neuronal projections

The structural framework of the reconstructed neurons consists of a series of nodes, each defined by spatial coordinates and interconnected by line segments. With reference to the CCFv3 atlas, the terminal nodes at the endpoints of axonal fibers (axon terminals) in the registered reconstructed neurons were quantified to determine the distribution of axonal terminals across different regions. The number of axonal terminals for each reconstructed neuron in each brain region was defined as its projection strength (PS) in that region, with the total PS across all regions constituting the projectome of each neuron, and the analysis performed in MATLAB. Hierarchical clustering based on the projectome of each reconstructed neuron was performed using the ’hclust’ function in R software with the Ward aggregation method.

#### General quantification and statistics

All the statistical details including the statistical methods, sample numbers, and *P* values can be found in the figures and/or figure legends. All data collection and data analyses were blinded in this study. GraphPad Prism (version 10.0), R software (version 4.1.2), and Microsoft Excel were used for statistical analyses. Group data are presented as bar plots showing mean ± standard error of the mean (SEM). Two-tailed unpaired Student’s *t* test, one-way analysis of variance (ANOVA), two-way ANOVA, and analysis of covariance (ANCOVA) were used to quantify the performance. The statistical significance was indicated as follows: *, *P* < 0.05; **, *P* < 0.01; ***, *P* < 0.001; N.S., not significant.

---

**Figure S1.**
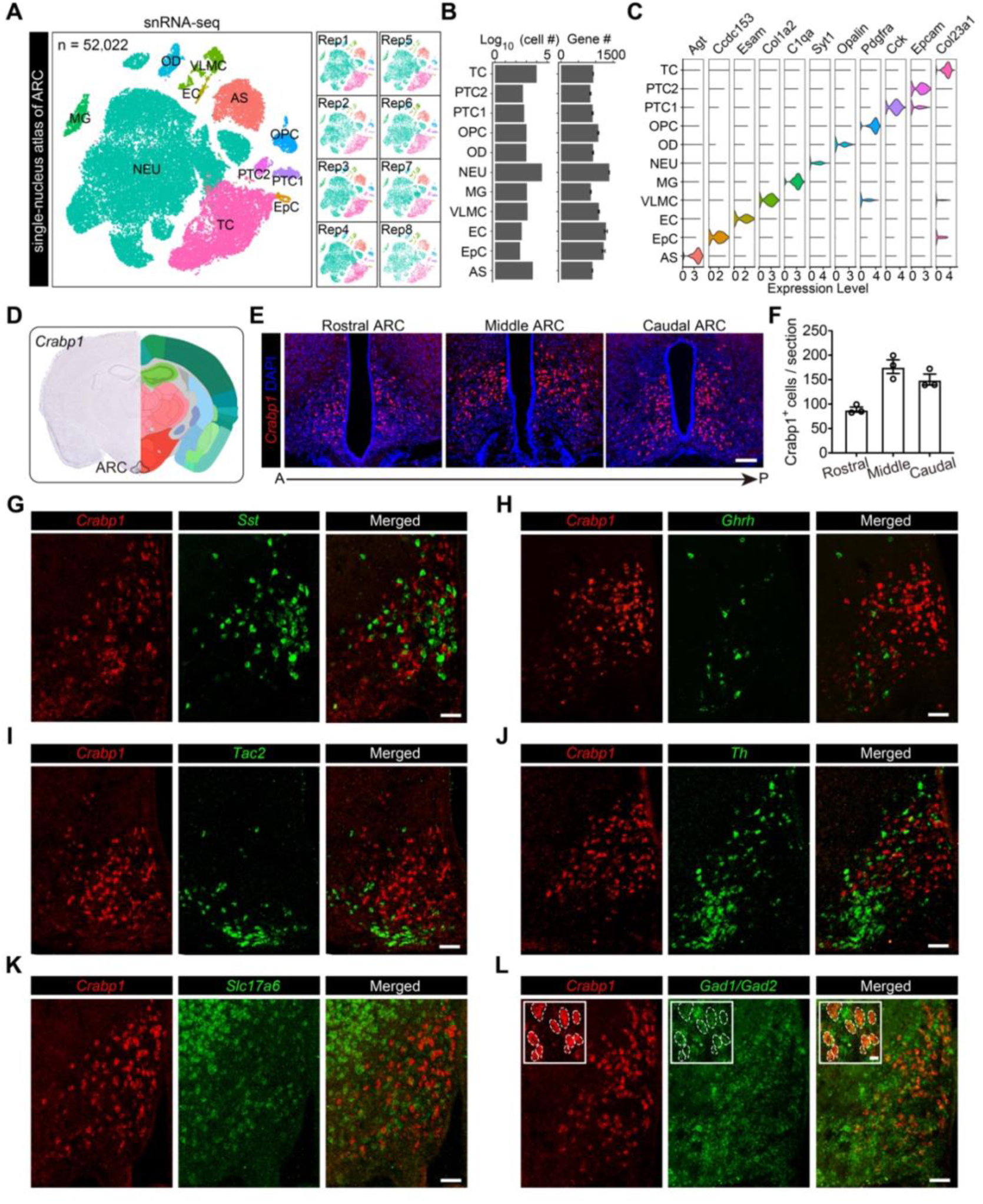
Spatial distribution of Crabp1 neurons that are GABAergic and distinct from *Sst*-, *Ghrh*-, *Tac2*-, and *Th*-expressing neurons, related to Figure 1. (**A**) Clustering analysis of 52,022 nuclei obtained from the arcuate hypothalamus. The arcuate nucleus (ARC) was microdissected and dissociated for single-nucleus RNA sequencing (snRNA-seq; n = 8 replicates). NEU, neuron; OPC, oligodendrocyte precursor cell; OD, oligodendrocyte; AS, astrocyte; TC, tanycyte; EC, ependymal cell; MG, microglia; VLMC, vascular and leptomeningeal cells; EpC, epithelial cell; PTC, pars tuberalis cells. (**B**) Bar plots showing the number of cells (left) and the average number of genes (right) for each cell type. (**C**) Violin plots displaying the expression levels of marker genes for each cell type. (**D**) Image showing the specific expression of *Crabp1* in the arcuate hypothalamus. *In situ* hybridization (ISH) of *Crabp1* mRNA was sourced from the Allen Mouse Brain Atlas and aligned with a reference atlas. (**E**) Representative confocal images showing the widespread distribution of *Crabp1*, spanning from anterior to posterior ARC. The sections were subjected to single molecule fluorescence *in situ* hybridization (smFISH) for detecting *Crabp1* mRNA and counterstained with DAPI (n = 3 brains). A, anterior; P, posterior. Scale bar, 50 μm. (**F**) Quantification of *Crabp1*-expressing cells across from anterior to posterior ARC (n = 23 sections from each of 3 brains). (**G**-**L**) Representative images showing that *Crabp1* mRNA do not co-localize with *Sst*, *Ghrh*, *Tac2*, *Th*, *Slc17a6*, but are co-stained with *Gad1/Gad2* in the ARC. Dual-color smFISH was used to assess the co-localization of *Crabp1* with markers of various arcuate neuronal subtypes. Scale bars, 50 μm.

**Figure S2.**
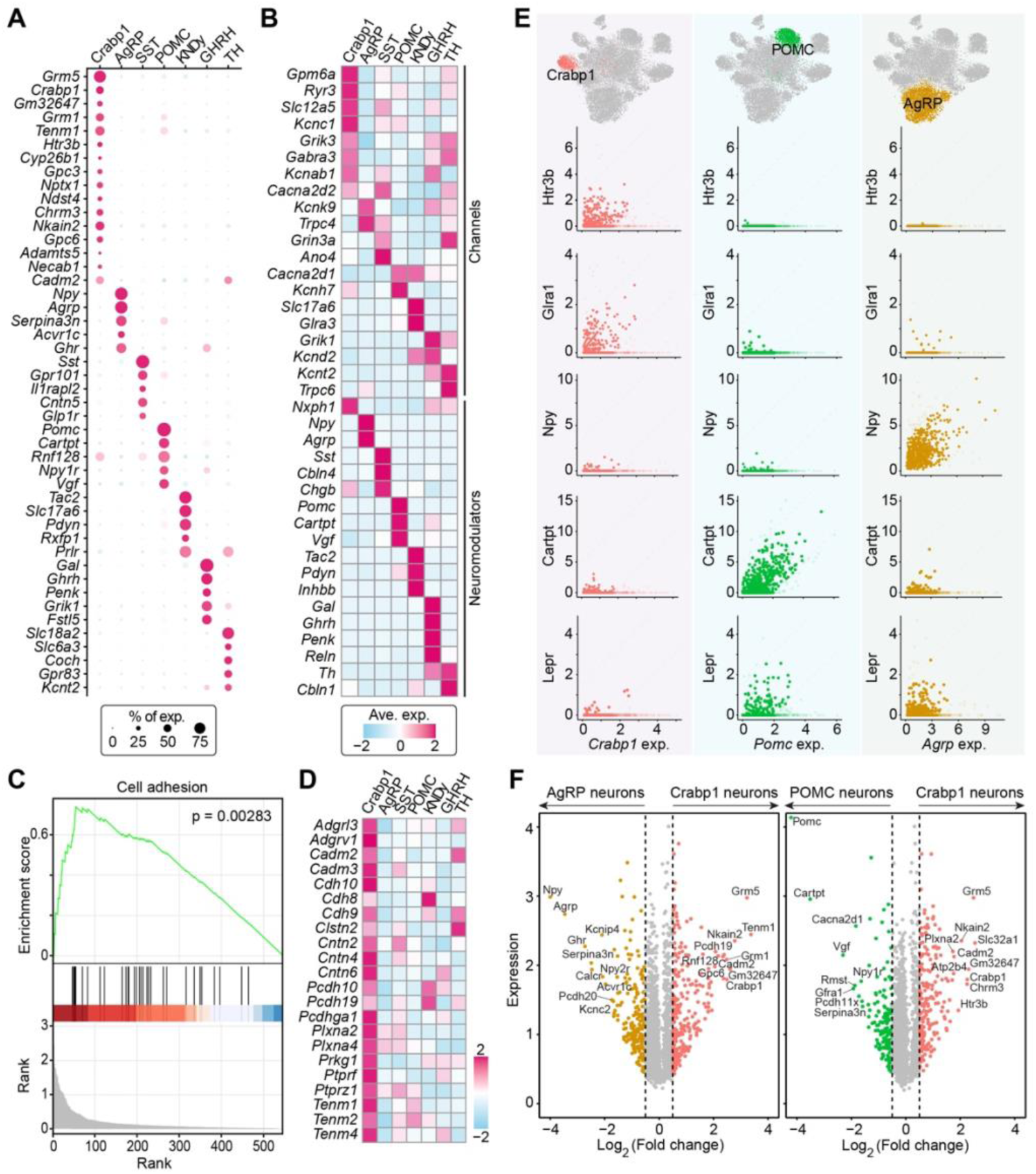
Molecular characterization of Crabp1 neurons, related to Figure 1. (**A**) Dot plot showing the expression of top marker genes specific to distinct neuronal subtypes in the ARC. (**B**) Heatmap showing the enriched expression of genes encoding channels and neuromodulators in various arcuate neuronal subtypes. (**C**) Gene set enrichment analysis (GSEA) indicating the abundance of genes associated with cell adhesion in Crabp1 neurons. (**D**) Heatmap showing the top cell adhesion related genes enriched in Crabp1 neurons. (**E**) Scatter plots showing the strong correlation between *Crabp1* and neurotransmitter receptor genes (e.g., *Htr3b* and *Glra1*), but not neuropeptides (e.g., *Npy* and *Cartpt*) or hormone receptors (e.g., *Lepr*). UMAP visualizations of Crabp1, AgRP, and POMC neurons are shown on the top. Gene expression levels are presented as counts per million mapped reads. (**F**) Volcano plots indicating differentially expressed genes between Crabp1 neurons and AgRP or POMC neurons. Genes with adjusted *P* value < 0.05 and fold change > 1.4 are highlighted.

**Figure S3.**
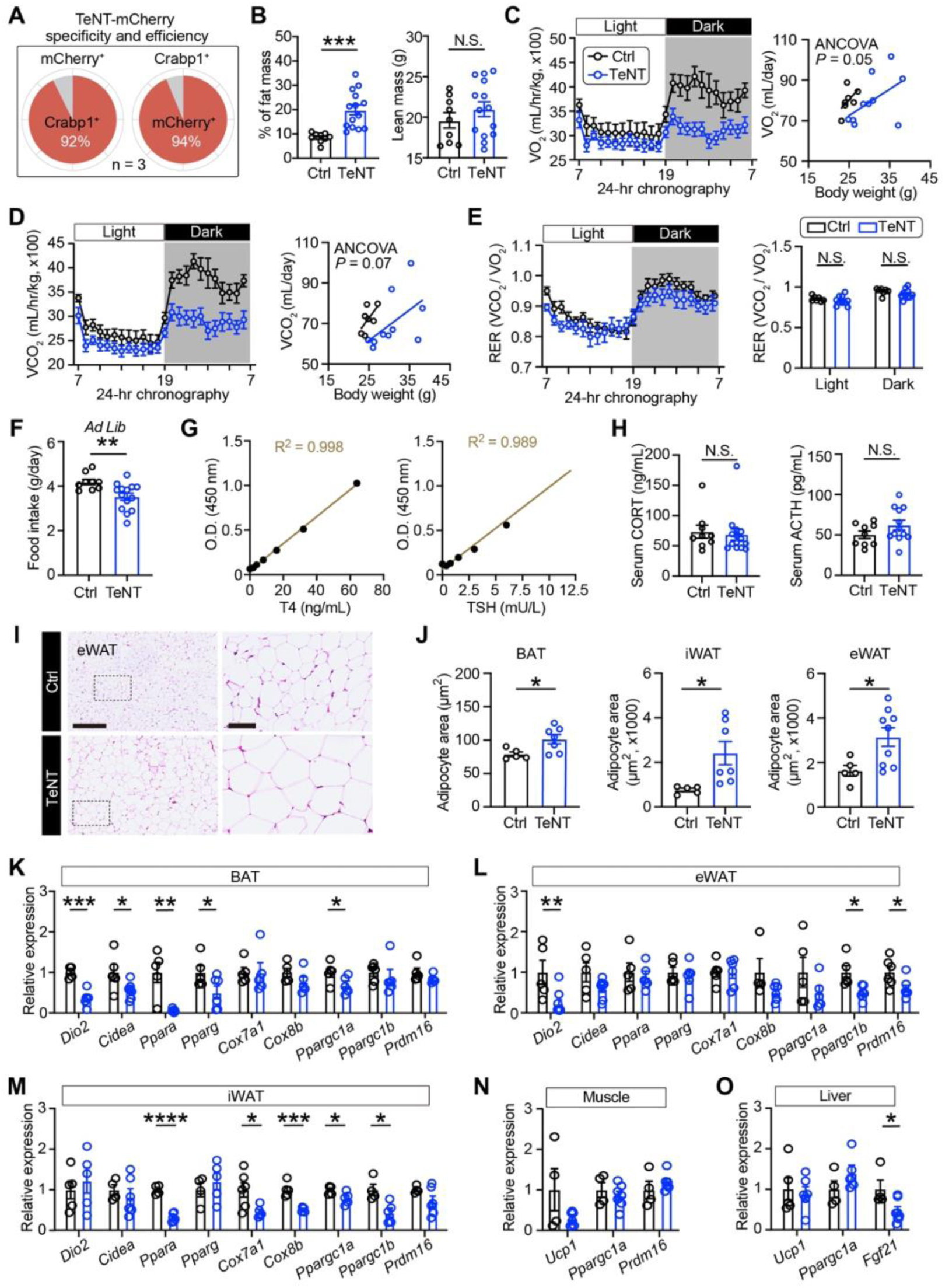
Disrupted synaptic transmission of Crabp1 neurons impairs energy metabolism, related to Figure 2. (**A**) Quantitative analysis showing the AAV-mediated labeling specificity and efficiency in arcuate Crabp1 neurons with *Crabp1-iCre* mice (n = 3). (**B**) Body fat percentage (left) and lean mass (right) in control mice and Crabp1 neuron-inactivated mice after 4 weeks of manipulation (n = 9 for control group; n = 14 for TeNT group). (**C** and **D**) Oxygen (O_2_) consumption (C) and carbon dioxide (CO_2_) production (D), along with regression analysis against body weight for both control and Crabp1 neuron-silenced mice (n = 7 for control group; n = 10 for TeNT group). Statistical analysis was performed using regression-based analysis of covariance (ANCOVA) with O_2_ or CO_2_ as dependent variable and body mass as covariate. (**E**) Mean curves (left) and bar plots (right) showing respiratory exchange ratio (RER) in *ad libitum* chow-fed mice during light and dark phases (n = 7 for control group; n = 10 for TeNT group). (**F**) Quantification of daily food intake under *ad libitum* condition (n = 9 for control group; n = 14 for TeNT group). (**G**) Standard curves employed to compute serum tetraiodothyronine (T4) and thyroid-stimulating hormone (TSH) levels. The measured optical density (OD) was linearly proportional to serum T4 and TSH levels. (**H**) Quantification of serum cortisol (CORT), and adrenocorticotropic hormone (ACTH) levels (n = 9 for control group; n = 12 for TeNT group). (**I**) Sample hematoxylin and eosin (H&E) staining images of epididymal white adipose tissue (eWAT), with magnified boxed regions highlighted. Scale bars, 250 μm and 50 μm for magnified images. (**J**) Quantitative analysis of average adipocyte area in brown adipose tissue (BAT), inguinal white adipose tissue (iWAT), and eWAT (n = 5 for control group; n = 7∼9 for TeNT group). (**K**-**O**) Thermogenesis-related gene expression in brown adipose tissue (BAT) (K), eWAT (L), iWAT (M), muscle (N), and liver (O) (n = 5∼6 for control group; n = 5∼8 for TeNT group). Values are shown as mean ± SEM. Significance was analyzed by unpaired two-tailed Student’s *t* test, unless otherwise denoted. N.S., not significant; *, *P* < 0.05; **, *P* < 0.01; ***, *P* < 0.001.

**Figure S4.**
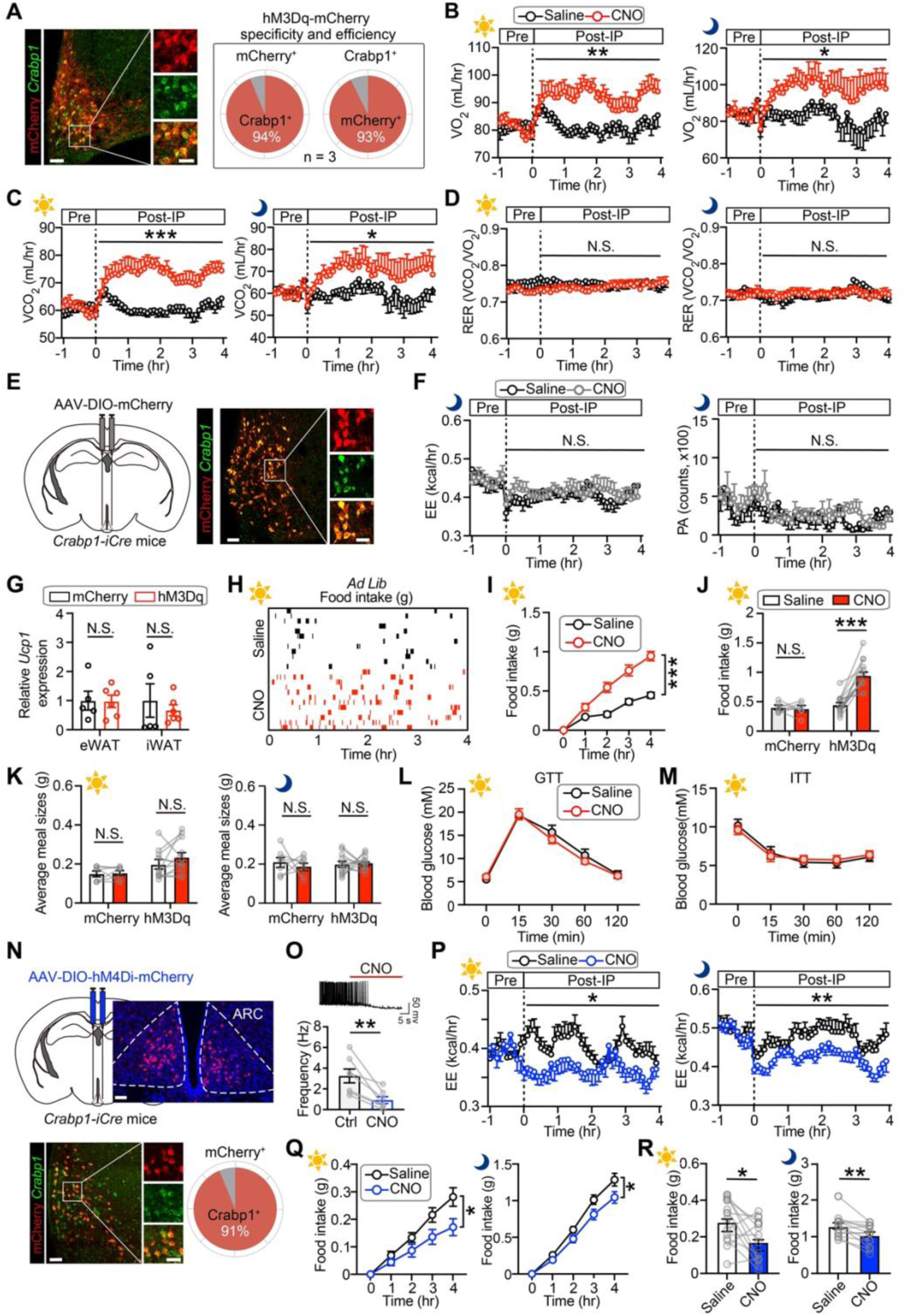
Chemogenetic manipulation of hypothalamic Crabp1 neurons regulates energy metabolism, related to Figure 3. (**A**) Specificity and efficiency of hM3Dq expression in Crabp1 neurons. Sample images showing the colocalization of Cre-driven mCherry expression with endogenous *Crabp1* mRNA puncta (left), alongside with quantitative analysis (right; n = 3 brains), are presented. Scale bars, 50 μm and 25 μm for magnified images. (**B**-**D**) Mean curves showing real-time O2 consumption (B), CO2 production (C) and RER (D) before (Pre-IP) and after (Post-IP) intraperitoneal injection of clozapine N-oxide (CNO; n = 8∼11 mice) or saline (n = 8∼11 mice) during light (left) and dark (right) phases. (**E**) Histological validation of mCherry expression in Crabp1 neurons. Scale bars, 50 μm and 25 μm for magnified images. (**F**) Mean curves showing real-time EE (left) and locomotion activity (right) before and after intraperitoneal injection of CNO or saline during dark phases (n = 10). (**G**) Gene expression analysis of *Ucp1* levels in eWAT and iWAT (n = 5 for saline group; n = 6 for CNO-activated group). (**H** and **I**) Raster plots (H) and mean curves (I) showing the dynamics of food consumption in *ad libitum*-fed mice within a 4 hours observation window following CNO or saline injection during the light cycle. These mice expressed hM3Dq in Crabp1 neurons (n = 12∼15). (**J**) Quantification of 4 hours food intake after CNO or saline administration during the light cycle. Control mice were microinjected with AAV-DIO-mCherry, while the experimental group were injected with AAV-DIO-hM3Dq-mCherry (n = 7 for control group; n = 15 for hM3Dq group). (**K**) Quantitative analysis of average meal size in control (n = 7) and hM3Dq-activated group (n = 15) during light (left) and dark (right) cycles. (**L** and **M**) Time course of blood glucose levels during the glucose tolerance test (GTT, L) and insulin tolerance test (ITT, M) at 4 hours post CNO (n = 9) or saline (n = 9) injection. (**N**) Specificity of hM4Di expression in Crabp1 neurons. Top: Expression of hM4Di-mCherry in the ARC. Scale bar, 50 μm. Bottom: Sample images (left) and quantitative analysis (right) of hM4Di-mCherry expression colocalizing with *Crabp1* mRNA puncta (n = 3 brains). Scale bars, 50 and 25 μm. (**O**) Electrophysiological validation of chemogenetic inhibition of Crabp1 neurons. Sample electrophysiological trace (top) and quantification (bottom) of action potential firing frequency in Crabp1 neurons before and after CNO perfusion on brain slices (n = 7 cells). (**P**) Mean curves showing real-time EE before and after CNO or saline administration during light (left) and dark (right) phases (n = 10 mice). (**Q**) Dynamics of food consumption in *ad libitum*-fed mice within a 4 hours period following CNO or saline injection during the light (left) or dark cycle (right). (**R**) Quantification of 4 hours food intake during the light (left) or dark cycle (right). Values are presented as mean ± SEM. Significance was assessed using paired or unpaired two-tailed Student’s *t* test and two-way analysis of variance (ANOVA) with Dunnett’s multiple comparison test. N.S., not significant; *, *P* < 0.05; **, *P* < 0.01; ***, *P* < 0.001.

**Figure S5.**
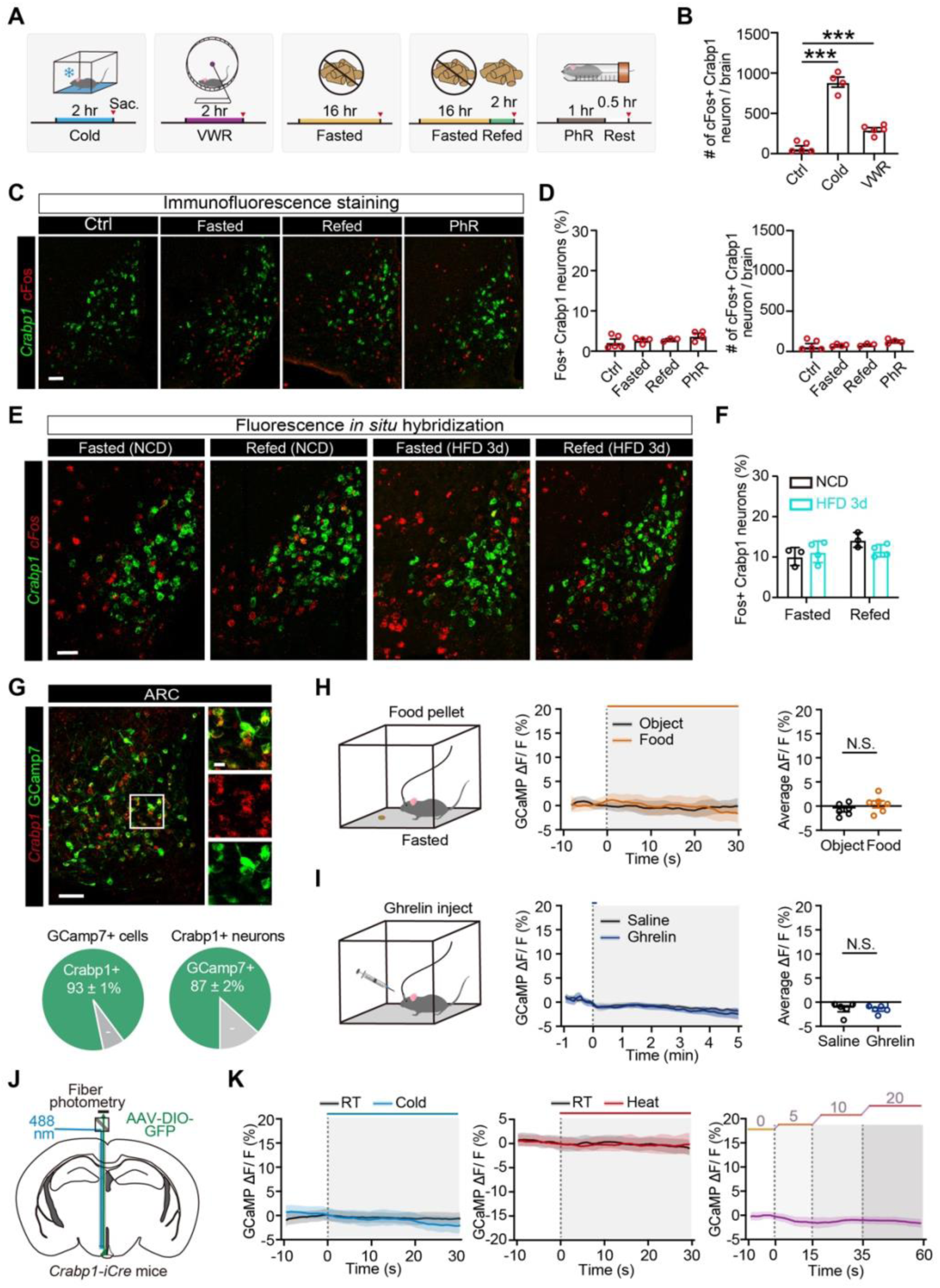
Crabp1 neurons fail to respond to hunger, satiety, and physical restraint, related to Figure 4. (**A**) Experimental scheme for detecting cFos expression in the ARC following the exposure to five different environmental or internal stimuli. VWR, voluntary wheel running; PhR, physical restraint; Sac, sacrifice. (**B**) Quantification of the cell number of cFos^+^ Crabp1 neurons following cold exposure or VWR (n = 4∼6). (**C** and **D**) Sample confocal images and quantification of the cFos immunoreactivity and Crabp1 mRNA puncta in the ARC following challenges of fasting, refeeding, and PhR under normal chow condition (n = 3∼5). Scale bar, 50 μm. (**E** and **F**) Representative confocal images and quantification of the dual-colour smFISH of *Fos* and *Crabp1* in the ARC following challenges of fasting and refeeding under high-fat diet condition (n = 3∼4). Scale bar, 50 μm. (**G**) Representative images (top) and quantitative analysis (bottom) of Cre-driven GCaMP7s expression colocalizing with endogenous *Crabp1* mRNA puncta (n = 3 brains). Scale bars, 50 μm and 35 μm for magnified images. (**H**) Neural dynamics of Crabp1 neurons upon food intake of a small chow pellet and a non-food object following overnight fasting. Experimental scheme (left) and mean calcium traces (middle) illustrating the response of Crabp1 neurons to object (black) and food (orange). The change in GCaMP fluorescence (ΔF/F) reflects neuronal activity. Scattered bar plots (right) showing the average calcium response of Crabp1 neurons to object or food (n = 6). (I) Neural dynamics of Crabp1 neurons following ghrelin or saline injection. Experimental scheme (left) and mean calcium traces (middle) showing the response of Crabp1 neurons to saline (black) or ghrelin (blue). Scattered bar plots (right) showing the average calcium response of Crabp1 neurons to saline or ghrelin (n = 5). Values represent mean ± SEM. (**J**) Viral transduction with AAV-DIO-EGFP as a control and surgical implantation of optical cannula. (**K**) Fluorescent dynamics in Crabp1 neurons expressing EGFP under cold exposure (left), heat exposure (middle), and exercise treadmill (right). Significance was analyzed by one-way ANOVA with Dunnett’s multiple comparison test and unpaired or paired two-tailed Student’s *t* test. N.S., not significant; ***, *P* < 0.001.

**Figure S6.**
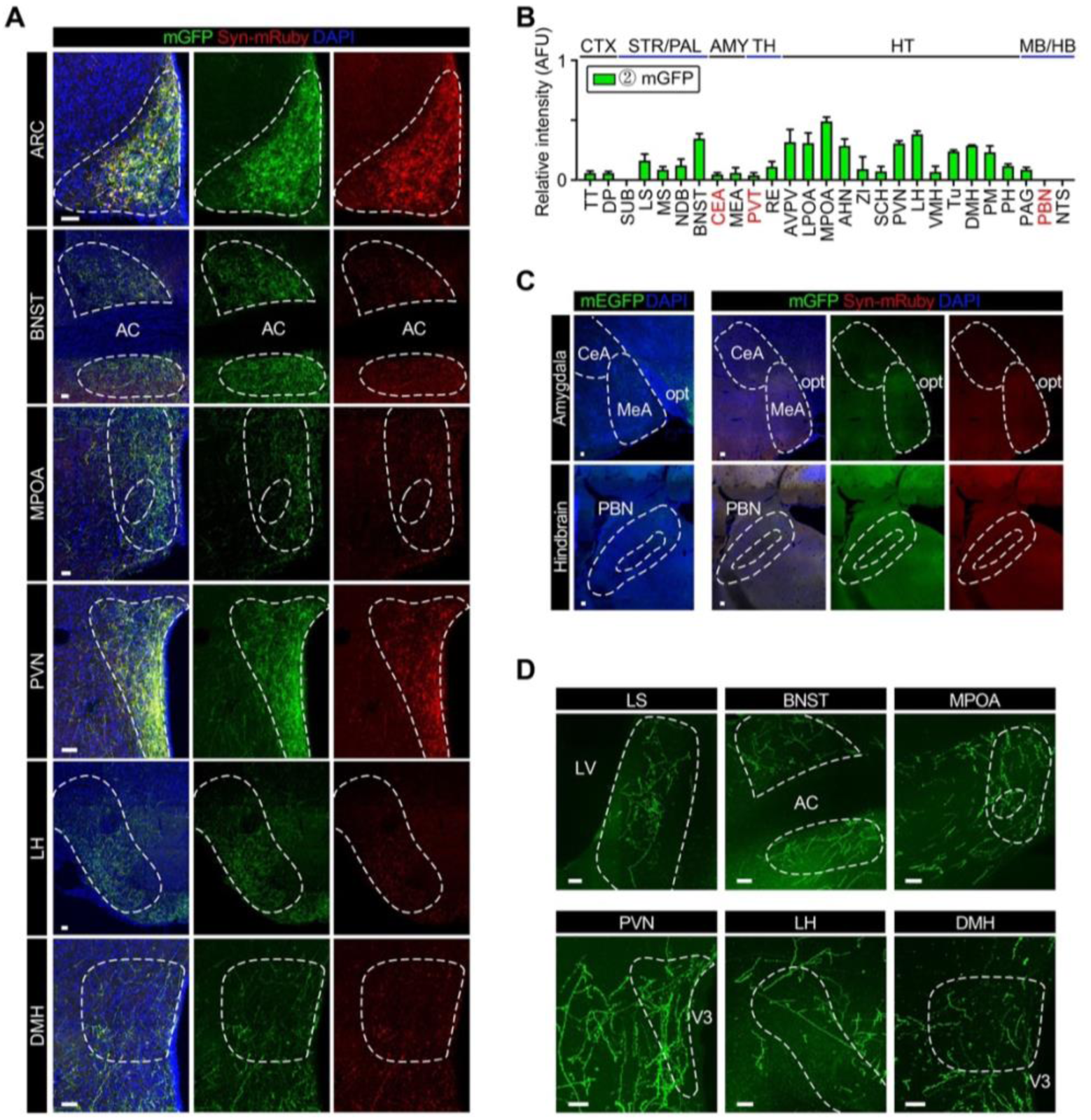
Structural connectivity of Crabp1 neurons at population and single-neuron levels, related to Figure 5. (**A**) Sample confocal images showing the genetic labeling of Crabp1 neurons in the ARC as well as mGFP^+^ axonal fibers and mRuby^+^ axonal boutons in multiple downstream target regions. See Table S6 for abbreviations. Scale bars, 50 μm. (**B**) Quantitative analysis of mGFP^+^ axonal fibers in various targeting brain regions (n = 3 brains). See Table S6 for abbreviations. Values represent mean ± SEM. (**C**) Representative images showing that the mEGFP^+^ axonal fibers (top) and mGFP^+^ axonal fibers along with mRuby^+^ axonal boutons (down) in the amygdala and hindbrain are barely detected. Scale bars, 50 μm. (**D**) Representative confocal images showing the fluorescence-labeled axonal fibers in multiple downstream target regions, which were labeled by AAV-CSSP-YFP and imaged using the fluorescence micro-optical sectioning tomography (fMOST) system. Scale bars, 50 μm.

**Figure S7.**
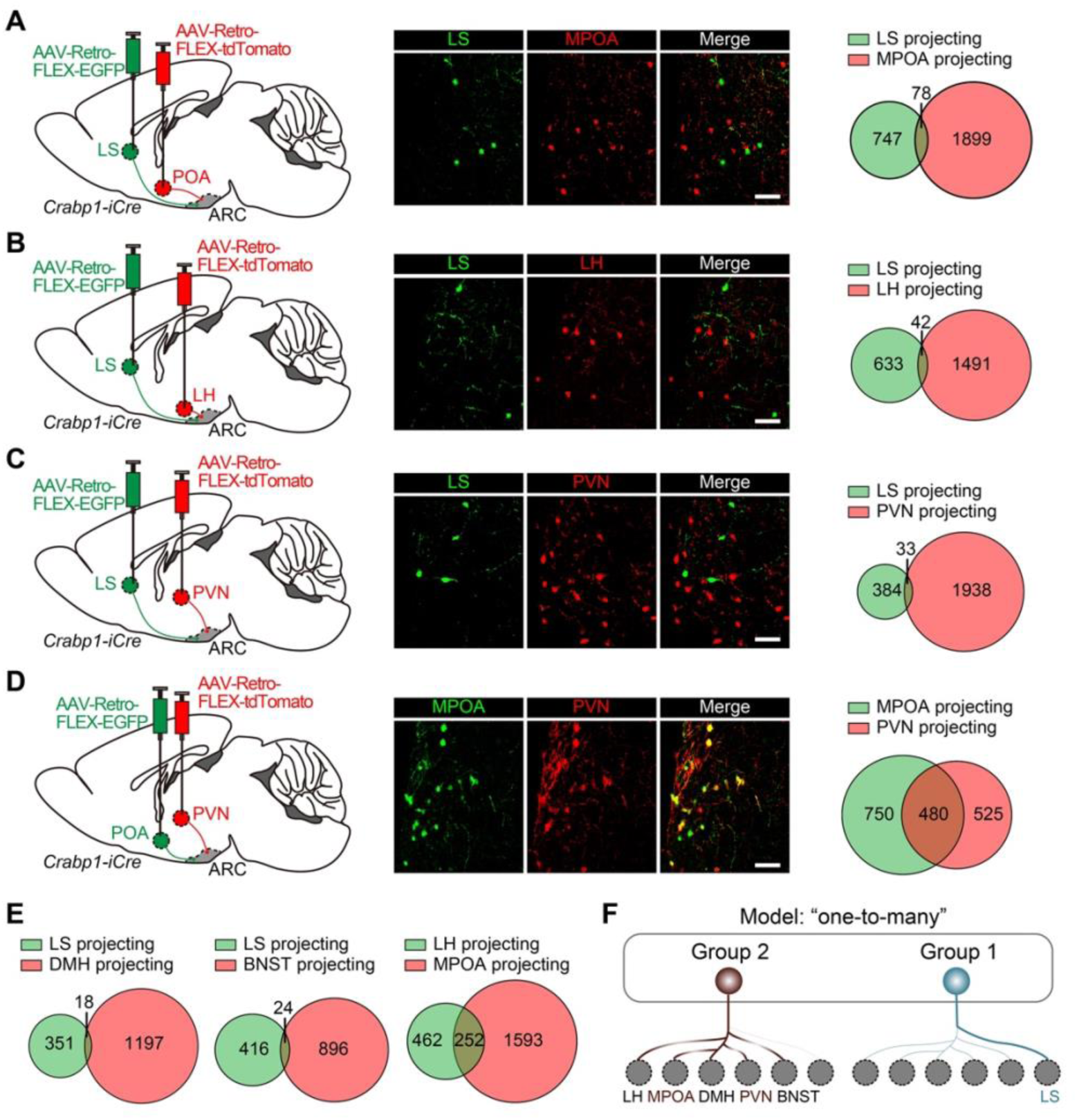
One-to-many projection pattern of Crabp1 neurons, related to Figure 5. (**A**-**C**) Minimal overlap between LS-projecting neurons and those targeting the MPOA, LH, and PVN. Schematics illustrating the experimental paradigms (left), representative images showing the overlapping between Crabp1 neurons targeting distinct regions (middle) and Venn diagrams quantifying the overlap (right; n = 3∼4 brains) are presented. Retrograde viruses, including AAV-Retro-FLEX-EGFP and AAV-Retro-FLEX-tdTomato, were injected into two distinct downstream projection regions in *Crabp1-iCre* mice. Scale bars, 50 μm. (**D**) Robust overlap between MPOA- and PVN-projecting Crabp1 neurons (n = 3∼4 brains). Scale bar, 50 μm. (**E**) Venn diagrams (n = 3∼4 brains for each pair) showing the overlap between LS- and DMH-projecting (left), LS- and BNST-projecting (middle), and LH- and MPOA-projecting Crabp1 neurons (right). (**F**) Schematic model summarizing the divergent axonal projection patterns between Group 1 and Group 2 neurons, supporting a “one-to-many” wiring configuration.

**Figure S8.**
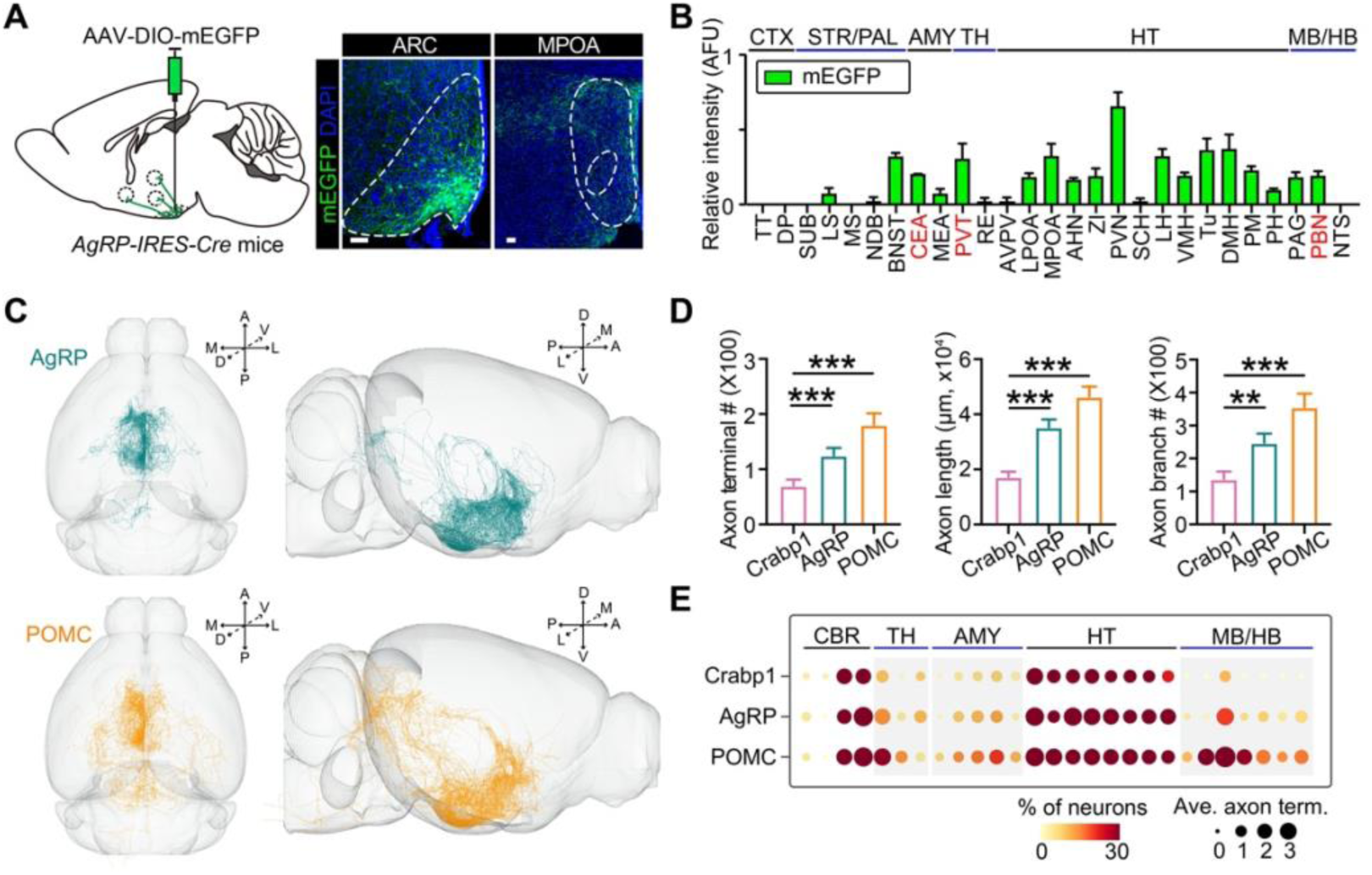
Comparison of structural connectivity between AgRP, POMC and Crabp1 neurons, related to Figure 5. (**A**) Population-level projection patterns of AgRP neurons. Left: Schematic diagram showing the viral labeling of AgRP neurons for anterograde tracing. Right: Representative images showing the genetic labeling of AgRP neurons and mEGFP^+^ axonal fibers in MPOA. Scale bars, 50 μm. (**B**) Quantitative analyses of the mEGFP^+^ axonal fibers of AgRP neurons in various targeting brain regions (n = 3 brains). (**C**) Three-dimensionally reconstructed AgRP and POMC neurons registered to the standard Allen Mouse Brain Atlas are shown in horizontal (left) and sagittal (right) views. A, anterior; P, posterior; D, dorsal; V, ventral; M, medial; L, lateral. (**D**) Quantitative analyses of axon terminal number, axon length, and axon branches of reconstructed AgRP, POMC and Crabp1 neurons. (**E**) Dot plot demonstrating the brain-wide projection patterns of AgRP, POMC, and Crabp1 neurons in each brain region. The dot size and intensity represent the percentage of neurons projecting to each brain area and the average number of terminals, respectively. Values represent mean ± SEM. Significance was analyzed by one-way ANOVA. **, *P* < 0.01; ***, *P* < 0.001.

**Figure S9.**
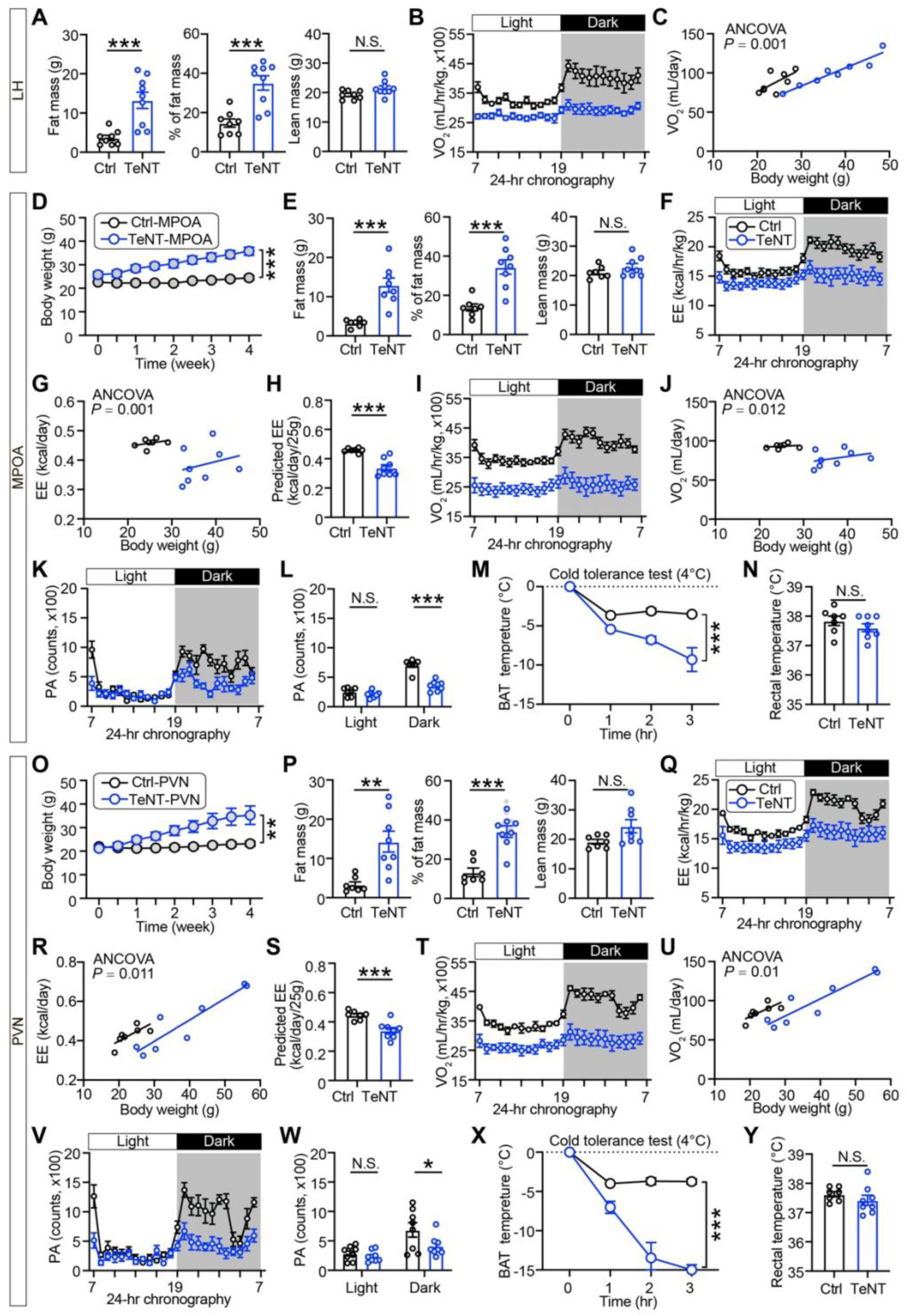
Metabolic disorders caused by inactivation of Crabp1 neurons that project to LH, MPOA, and PVN, related to Figure 6. (**A**-**C**) Body composition (A) and O_2_ consumption (B and C) in mice subjected to synaptic silencing of the ARC^Crabp1^-LH pathway (n = 8∼9). Animals receiving AAV-fDIO-mCherry injection in the ARC used as controls. (**D**-**N**) Body weight changes (D), body composition (E), total EE (F to H), O_2_ consumption (I and J), locomotion activity (K and L), adaptive BAT thermogenesis during cold exposure (M), and rectal temperature (N) in mice subjected to synaptic silencing of the ARC^Crabp1^-MPOA pathway (n = 7∼8). Animals injected with AAV-fDIO-mCherry in the ARC were used as controls. (**O**-**Y**) Body weight changes (O), body composition (P), total EE (Q to S), O_2_ consumption (T and U), locomotion activity (V and W), adaptive BAT thermogenesis during cold exposure (X), and rectal temperature (Y) in mice subjected to synaptic silencing of the ARC^Crabp1^-PVN pathway (n = 7∼8). The control group received AAV-fDIO-mCherry injection in the ARC. Values are presented as mean ± SEM. Statistical significance was analyzed by two-way ANOVA with Sidak’s multiple comparison test or unpaired two-tailed Student’s *t* test. Statistical analysis was performed using regression-based ANCOVA with EE or O_2_ as dependent variable and body mass as covariate. N.S., not significant; *, *P* < 0.05; **, *P* < 0.01; ***, *P* < 0.001.

**Figure S10.**
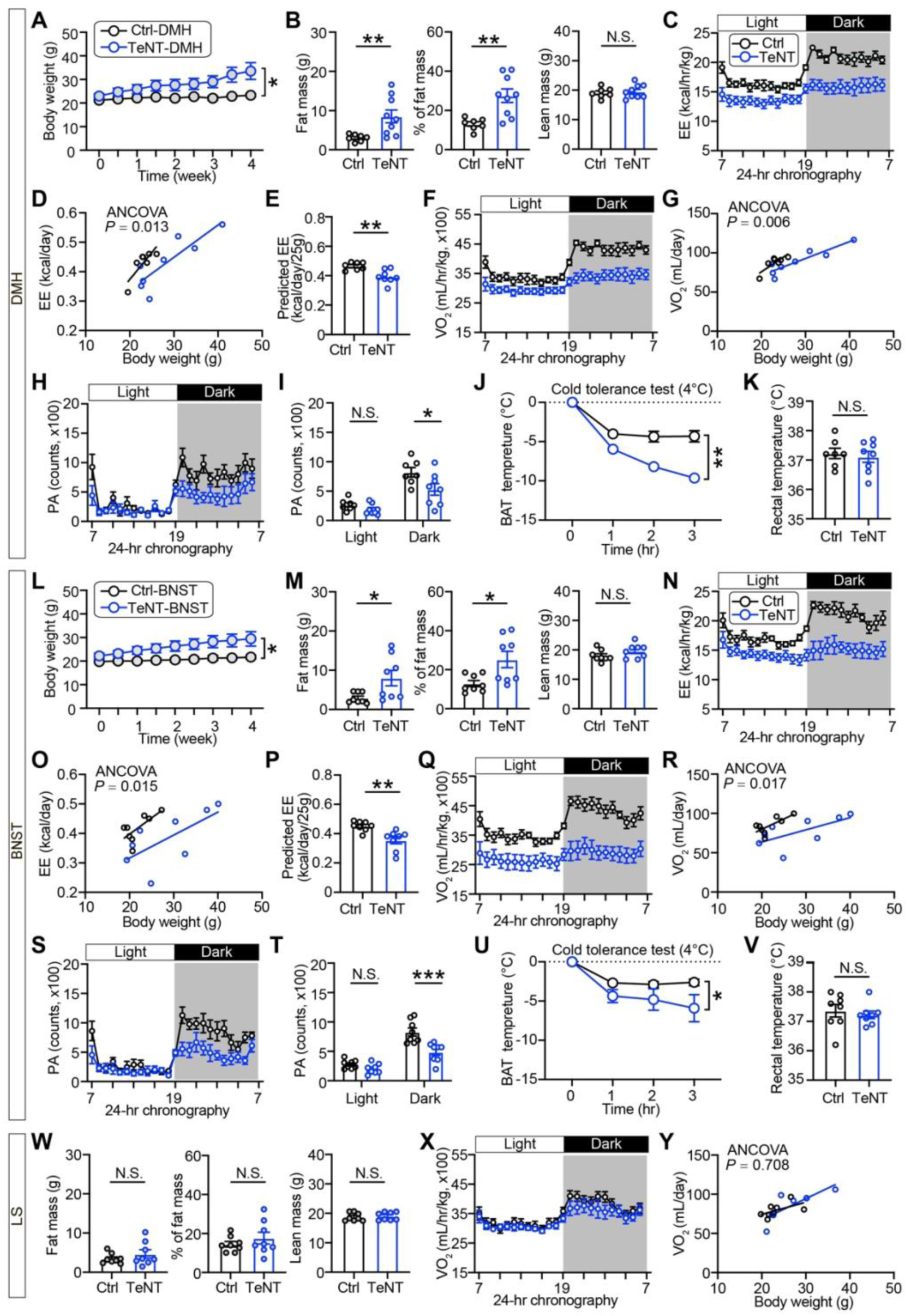
Metabolic disorders caused by inactivation of Crabp1 neurons that project to DMH, BNST, but not LS, related to Figure 6. (**A**-**K**) Body weight changes (A), body composition (B), total EE (C to E), O_2_ consumption (F and G), locomotion activity (H and I), adaptive BAT thermogenesis during cold exposure (J), and rectal temperature (K) in mice subjected to synaptic silencing of the ARC^Crabp1^-DMH pathway (n = 7∼8). Animals injected with AAV-fDIO-mCherry in the ARC were used as controls. (**L**-**V**) Body weight changes (L), body composition (M), total EE (N to P), O_2_ consumption (Q and R), locomotion activity (S and T), adaptive BAT thermogenesis during cold exposure (U), and rectal temperature (V) in mice subjected to synaptic silencing of the ARC^Crabp1^-BNST pathway (n = 7∼8). The control group received AAV-fDIO-mCherry injection in the ARC. (**W**-**Y**) Body composition (W) and O_2_ consumption (X and Y) in mice subjected to synaptic silencing of the ARC^Crabp1^-LS pathway (n = 8∼9). Animals receiving AAV-fDIO-mCherry injection in the ARC served as controls. Values are presented as mean ± SEM. Statistical significance was analyzed by two-way ANOVA with Sidak’s multiple comparison test or unpaired two-tailed Student’s *t* test. Statistical analysis was performed using regression-based ANCOVA with EE or O_2_ as dependent variable and body mass as covariate. N.S., not significant; *, *P* < 0.05; **, *P* < 0.01; ***, *P* < 0.001.

**Figure S11.**
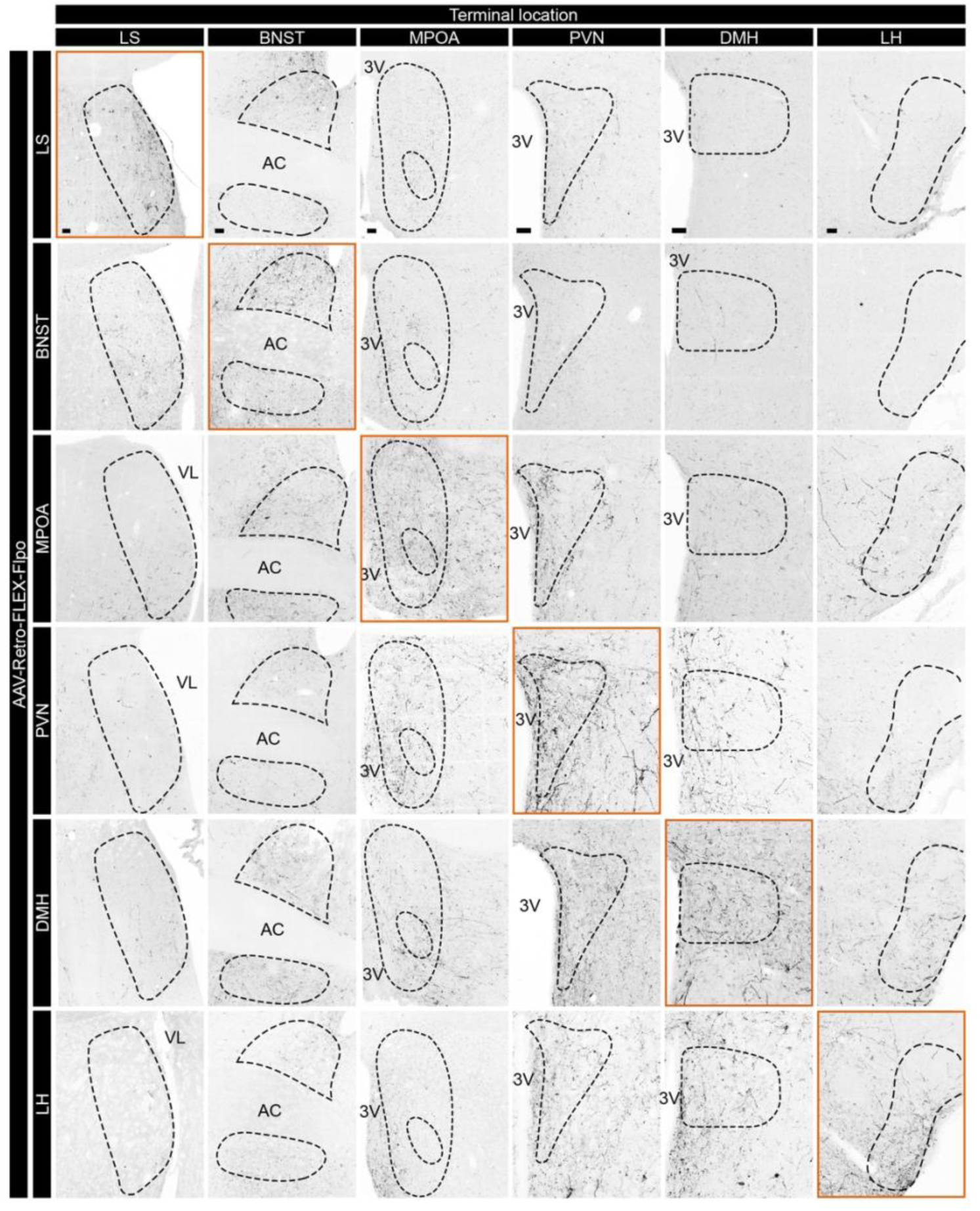
Axon collateral branching pattern of Crabp1 neurons, related to Figure 6. Axon fibers were detected in six predominant downstream projection targets (denoted by column headings) of Crabp1 neurons following the injection of AAV-Retro-FLEX-Flpo into six efferent sites (denoted by row headings), alongside AAV encoding Flpo-dependent mCherry into the ARC in *Crabp1-iCre* mice. Orange boxes mark the retrograde AAV injection sites. Scale bars, 50 μm.

